# Molecular evolutionary trends and feeding ecology diversification in the Hemiptera, anchored by the milkweed bug genome

**DOI:** 10.1101/201731

**Authors:** Kristen A. Panfilio, Iris M. Vargas Jentzsch, Joshua B. Benoit, Deniz Erezyilmaz, Yuichiro Suzuki, Stefano Colella, Hugh M. Robertson, Monica F. Poelchau, Robert M. Waterhouse, Panagiotis Ioannidis, Matthew T. Weirauch, Daniel S.T. Hughes, Shwetha C. Murali, John H. Werren, Chris G.C. Jacobs, Elizabeth J. Duncan, David Armisén, Barbara M.I. Vreede, Patrice Baa-Puyoulet, Chloé S. Berger, Chun-che Chang, Hsu Chao, Mei-Ju M. Chen, Yen-Ta Chen, Christopher P. Childers, Ariel D. Chipman, Andrew G. Cridge, Antonin J.J. Crumière, Peter K. Dearden, Elise M. Didion, Huyen Dinh, HarshaVardhan Doddapaneni, Amanda Dolan, Shannon Dugan, Cassandra G. Extavour, Gérard Febvay, Markus Friedrich, Neta Ginzburg, Yi Han, Peter Heger, Christopher J. Holmes, Thorsten Horn, Yi-min Hsiao, Emily C. Jennings, J. Spencer Johnston, Tamsin E. Jones, Jeffery W. Jones, Abderrahman Khila, Stefan Koelzer, Viera Kovacova, Megan Leask, Sandra L. Lee, Chien-Yueh Lee, Mackenzie R. Lovegrove, Hsiao-ling Lu, Yong Lu, Patricia J. Moore, Monica C. Munoz-Torres, Donna M. Muzny, Subba R. Palli, Nicolas Parisot, Leslie Pick, Megan Porter, Jiaxin Qu, Peter N. Refki, Rose Richter, Rolando Rivera Pomar, Andrew J. Rosendale, Siegfried Roth, Lena Sachs, M. Emília Santos, Jan Seibert, Essia Sghaier, Jayendra N. Shukla, Richard J. Stancliffe, Olivia Tidswell, Lucila Traverso, Maurijn van der Zee, Séverine Viala, Kim C. Worley, Evgeny M. Zdobnov, Richard A. Gibbs, Stephen Richards

## Abstract

**Background:** The Hemiptera (aphids, cicadas, and true bugs) are a key insect order, with high diversity for feeding ecology and excellent experimental tractability for molecular genetics. Building upon recent sequencing of hemipteran pests such as phloem-feeding aphids and blood-feeding bed bugs, we present the genome sequence and comparative analyses centered on the milkweed bug *Oncopeltus fasciatus*, a seed feeder of the family Lygaeidae.

**Results:** The 926-Mb *Oncopeltus* genome is well represented by the current assembly and official gene set. We use our genomic and RNA-seq data not only to characterize the protein-coding gene repertoire and perform isoform-specific RNAi, but also to elucidate patterns of molecular evolution and physiology. We find ongoing, lineage-specific expansion and diversification of repressive C2H2 zinc finger proteins. The discovery of intron gain and turnover specific to the Hemiptera also prompted evaluation of lineage and genome size as predictors of gene structure evolution. Furthermore, we identify enzymatic gains and losses that correlate with feeding biology, particularly for reductions associated with derived, fluid-nutrition feeding.

**Conclusions:** With the milkweed bug, we now have a critical mass of sequenced species for a hemimetabolous insect order and close outgroup to the Holometabola, substantially improving the diversity of insect genomics. We thereby define commonalities among the Hemiptera and delve into how hemipteran genomes reflect distinct feeding ecologies. Given *Oncopeltus'*s strength as an experimental model, these new sequence resources bolster the foundation for molecular research and highlight technical considerations for the analysis of medium-sized invertebrate genomes.

## Background

The number of animals with sequenced genomes continues to increase dramatically, and there are now over 100 insect species with assembled and annotated genomes [1]. However, the majority belong to the Holometabola (*e.g.*, flies, beetles, wasps, butterflies), the group characterized by a biphasic life history with distinct larval and adult phases separated by dramatic metamorphosis during a pupal stage. The Holometabola represent only a fraction of the full morphological and ecological diversity across the Insecta: over half of all orders are hemimetabolous. Imbalance in genomic resources limits the exploration of this diversity, including the environmental and developmental requirements of a hemimetabolous life style with a progression of flightless nymphal (juvenile) instars. Addressing this paucity, we report comparative analyses based on genome sequencing of the large milkweed bug, *Oncopeltus fasciatus*, as a hemimetabolous representative of the larger diversity of insects.

*Oncopeltus* is a member of the Hemiptera, the most species-rich hemimetabolous order. Together with the Thysanoptera and, traditionally, the Psocodea, the Hemiptera form the hemipteroid assemblage (or Acercaria), a close outgroup to the Holometabola [2, 3]. All Hemiptera share the same piercing and sucking mouthpart anatomy [4], yet they have diversified to exploit food sources ranging from seeds and plant tissues (phytophagy) to phloem sap (mucivory) and vertebrate blood (hematophagy). For this reason, many hemipterans are agricultural pests or human disease vectors, and genome sequencing efforts to date have focused on these species (Fig. 1, [5]), including phloem-feeding aphids [6-8], psyllids [9], and planthoppers [10], and the hematophagous kissing bug, *Rhodnius prolixus* [11], a vector of Chagas disease, and bed bug, *Cimex lectularius* [12, 13]. Building on transcriptomic data, genome projects are also in progress for other pest species within the same infraorder as *Oncopeltus*, such as the stink bug *Halyomorpha halys* [14, 15].

**Fig. 1.**
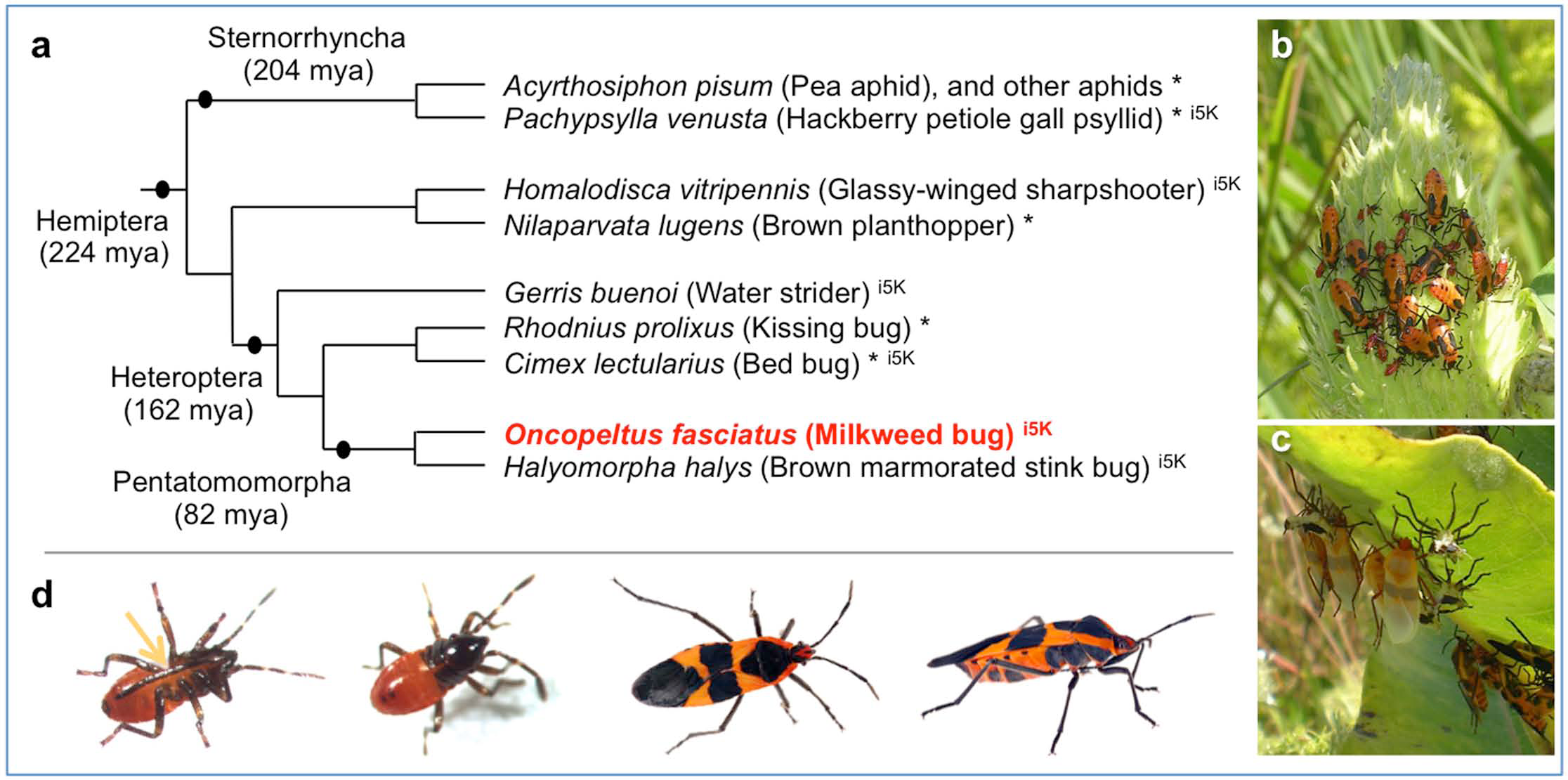
**The large milkweed bug, *Oncopeltus fasciatus*, shown in its phylogenetic and environmental context.** **(a)** Species tree of selected Hemiptera with genomic and transcriptomic resources, based on phylogenetic analyses and divergence time estimates in [3]. Species marked with an asterisk (*) have published resources; those with the appellation “i5K” are part of a current pilot project supported by the Baylor College of Medicine Human Genome Sequencing Center and the National Agricultural Library of the USDA. Note that recent analyses suggest the traditional infraorder Cimicomorpha, to which *Rhodnius* and *Cimex* belong, may be paraphyletic [16]. **(b-c)** Milkweed bugs on their native food source, the milkweed plant: gregarious nymphs of different instars on a milkweed seed pod (b), and pale, recently eclosed adults and their shed exuvia (c). Images were taken at Avalon Park and Preserve, Stony Brook, New York, USA, courtesy of Deniz Erezyilmaz, used with permission. **(d)** Individual bugs, shown from left to right: first instar nymphs (ventral and dorsal views) and adults (dorsal and lateral views); images courtesy of Kristen Panfilio (nymphs) and Jena Johnson (adults), used with permission. The arrow labels the labium (the “straw”), part of the hemipteran mouthpart anatomy adapted for feeding by piercing and sucking.

The milkweed bug has feeding ecology traits that are both conservative and complementary to those of previously sequenced hemipterans. Its phytophagy is ancestral for the large infraorder Pentatomomorpha and representative of most extant Hemiptera [16]. Moreover, as a seed feeder *Oncopeltus* has not undergone the marked life style changes associated with fluid feeding (mucivory or hematophagy), including dependence on endosymbiotic bacteria to provide nutrients lacking in the diet. Gene loss in the pea aphid, *Acyrthosiphon pisum*, makes it reliant on the obligate endosymbiont *Buchnera aphidicola* for synthesis of essential amino acids [6, 17]. Although hematophagy arose independently in *Rhodnius* and *Cimex* [16], their respective endosymbionts, *Rhodococcus rhodnii* and *Wolbachia*, must provide vitamins lacking in a blood diet [18]. In contrast, the seed-feeding subfamily Lygaeinae, including *Oncopeltus*, is notable for the absence of prominent endosymbiotic anatomy: these bugs lack both the midgut crypts that typically house bacteria and the bacteriomes and endosymbiotic balls seen in other Lygaeidae [19].

As the native food source of *Oncopeltus* is the toxic milkweed plant, its own feeding biology has a number of interesting implications regarding detoxification and sequestration of cardenolide compounds. A prominent consequence of this diet is the bright red-orange aposematic (warning) coloration seen in *Oncopeltus* embryos, nymphs, and adults [20, 21]. Thus, diet, metabolism, and body pigmentation are functionally linked biological features for which one may expect changes in gene repertoires to reflect diversity within an order, and the Hemiptera provide an excellent opportunity to explore this.

Furthermore, *Oncopeltus* has been an established laboratory model organism for over 60 years, with a rich experimental tradition in a wide range of studies from physiology and development to evolutionary ecology [21-23]. It is among the few experimentally tractable hemimetabolous insect species, and it is amenable to a range of molecular techniques (*e.g*., [24-26]). In fact, it was one of the first insect species to be functionally investigated by RNA interference (RNAi, [27]). RNAi in *Oncopeltus* is highly effective across different life history stages, which has led to a resurgence of experimental work over the past fifteen years, with a particular focus on the evolution of developmentally important regulatory genes (reviewed in [23]).

Here, we focus on these two themes – feeding biology diversity within the Hemiptera and *Oncopeltus* as a research model for macroevolutionary genetics. Key insights derive from a combination of global comparative genomics and detailed computational analyses that are supported by extensive manual curation, empirical data for gene expression, sequence validation, and new isoform-specific RNAi. We thereby identify genes with potentially restricted life history expression in *Oncopeltus* and that are unique to the Hemiptera, clarify evolutionary patterns of zinc finger protein expansion, categorize predictors of insect gene structure, and identify lateral gene transfer and amino acid metabolism features that correlate with feeding biology.

## Results and Discussion

### The genome and its assembly

*Oncopeltus fasciatus* has a diploid chromosome number (2*n*) of 16, comprised of seven autosomal pairs and two sex chromosomes with the XX/XY sex determination system [28, 29]. To analyze this genetic resource, we sequenced and assembled the genome using next-generation sequencing approaches (Table 1, see also Methods and Supplemental Notes Sections 1-4). We measure the genome size to be 923 Mb in females and 928 Mb in males based on flow cytometry data (Supplemental Note 2.1.a). The assembly thus contains 84% of the expected sequence, which is comparable to other recent, medium-sized insect genomes [12, 30]. However, our analyses of the *k*-mer frequency distribution in raw sequencing reads yielded ambiguous estimates of genome size and heterozygosity rate, which is suggestive of high heterozygosity and repetitive content ([31], Supplemental Note 2.1.b). In further analyses we indeed obtained high estimates of repetitive content, although heterozygosity does not unduly influence gene prediction (see below, based on protein orthology assessments). These computationally challenging features may be increasingly relevant as comparative genomics extends to insect species with larger genomes (>1 Gb) – a common feature among hemimetabolous insects [5, 32].

As template DNA was prepared from dissected adults from which gut material was removed, the resulting assembly is essentially free of contamination. Only five small scaffolds had high bacterial homology, each to a different, partial bacterial genome (Supplemental Note 2.2).

**Table 1.**
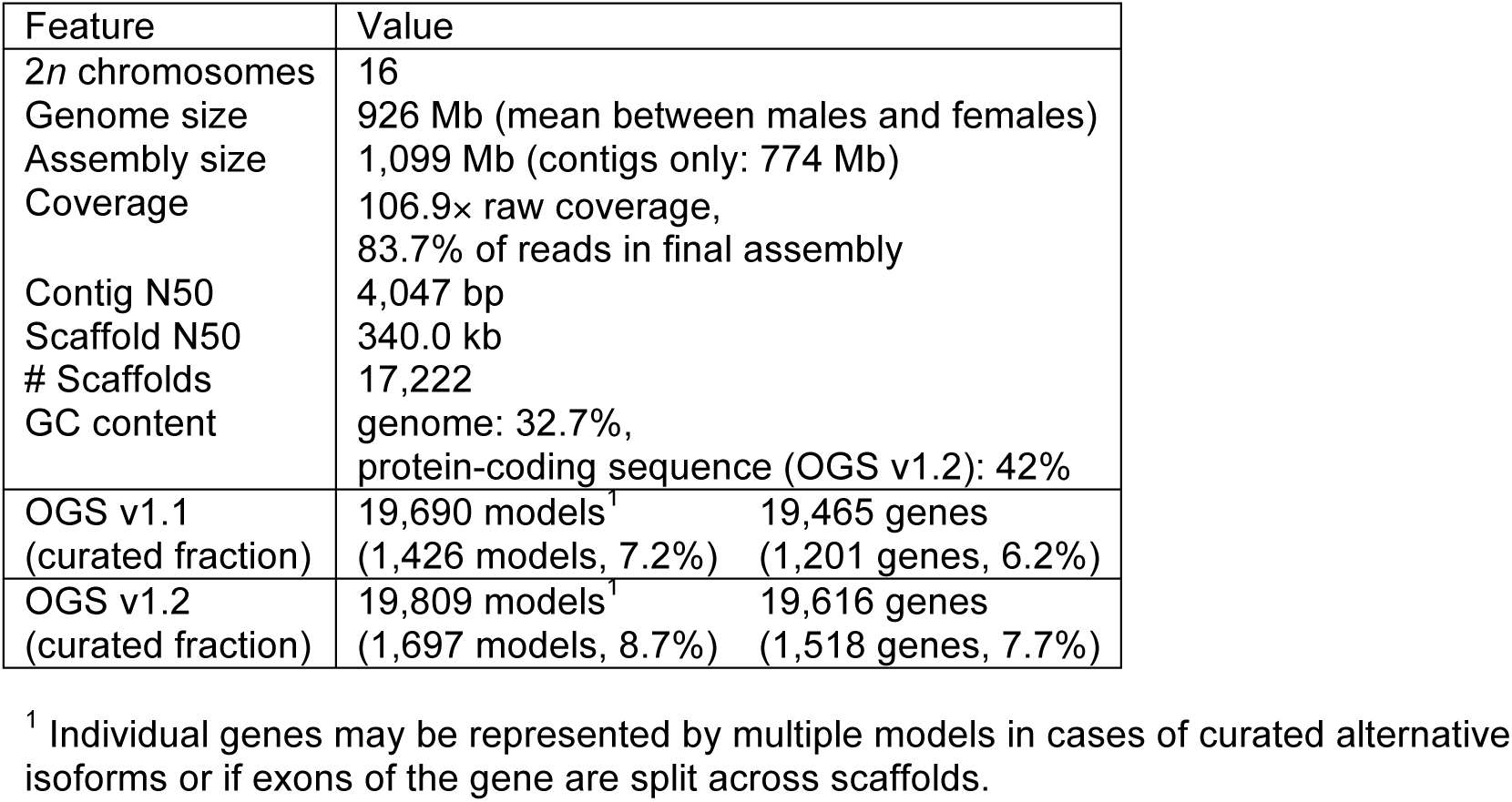
***Oncopeltus fasciatus* genome metrics.**

| Feature | Value |
| --- | --- |
| 2n chromosomes | 16 |
| Genome size | 926 Mb (mean between males and females) |
| Assembly size | 1,099 Mb (contigs only: 774 Mb) |
| Coverage | 106.9× raw coverage,<br>83.7% of reads in final assembly |
| Contig N50 | 4,047 bp |
| Scaffold N50 | 340.0 kb |
| # Scaffolds | 17,222 |
| GC content | genome: 32.7%,<br>protein-coding sequence (OGS v1.2): 42% |
| OGS v1.1<br>(curated fraction) | 19,690 models<sup>1</sup> 19,465 genes<br>(1,426 models, 7.2%) (1,201 genes, 6.2%) |
| OGS v1.2<br>(curated fraction) | 19,809 models<sup>1</sup> 19,616 genes<br>(1,697 models, 8.7%) (1,518 genes, 7.7%) |

<sup>1</sup> Individual genes may be represented by multiple models in cases of curated alternative isoforms or if exons of the gene are split across scaffolds.

### The official gene set and conserved gene linkage

The official gene set (OGS) was generated by automatic annotation followed by manual curation in a large-scale effort by the research community (Supplemental Notes Sections 3-4). Curation revised automatic models, added alternative isoforms and *de novo* models, and documented multiple models for genes split across scaffolds. We found that automatic predictions were rather conservative for hemipteran gene structure (see below). Thus, manual curation often extended gene loci as exons were added, including merging discrete automatic models (Supplemental Note 4, Table S4.4). The OGS v1.1 was generated for global analyses to characterize the gene repertoire. The latest version, OGS v1.2, primarily adds chemoreceptor genes of the ionotropic and odorant receptor classes and genes encoding metabolic enzymes. Altogether, the research community curated 1,697 gene models (8.7% of OGS v1.2), including 316 *de novo* models (Table S4.1, Supplemental Notes Section 5). The majority of curated models are for genes encoding cuticular proteins (11%), chemoreceptors (19%), and developmental regulators such as transcription factors and signaling pathway components (40%, including the BMP/TGF-β, Toll/NF-κB, Notch, Hedgehog, Torso RTK, and Wnt pathways).

In addition to assessing gene model quality, manual curation of genes whose orthologs are expected to occur in syntenic clusters also validates assembly scaffolding. Complete loci could be found for single orthologs of all Hox cluster genes, where *Hox3/zen* and *Hox4/Dfd* are linked in the current assembly and have ≥99.9% nucleotide identity with experimentally validated sequences ([33-35], Supplemental Note 5.1.b). Conserved linkage was also confirmed for the homeobox genes of the Iroquois complex, the Wnt ligands *wingless* and *wnt10*, and two linked pairs from the Runt transcription factor complex (Supplemental Notes 5.1.a, 5.1.c, 5.1.i, 5.1.j). Further evidence for correct scaffold assembly comes from the curation of large, multi-exonic loci. For example, the cell polarity and cytoskeletal regulator encoded by the conserved *furry* gene includes 47 exons spanning a 437-kb locus, which were all correctly assembled on a single scaffold.

### Gene expression profiles across the milkweed bug life cycle

To augment published transcriptomic resources [36, 37], we sequenced three different post-embryonic samples (“i5K” dataset, see Methods). We then compared the OGS to the resulting *de novo* transcriptome and to a previously published embryonic and maternal (ovary) transcriptome (“454” pyrosequencing dataset, [36]). Our OGS is quite comprehensive, containing 90% of transcripts from each transcriptomic dataset (Fig. 2a). The OGS also contains an additional 3,146 models (16% of OGS) not represented in either transcriptome, including 163 *de novo* models encoding chemoreceptors. Such genes are known for lineage-specific expansions and highly tissue- and stage-specific expression ([38, 39], and see below), and our OGS captures these genes with rare transcripts.

**Fig. 2.**
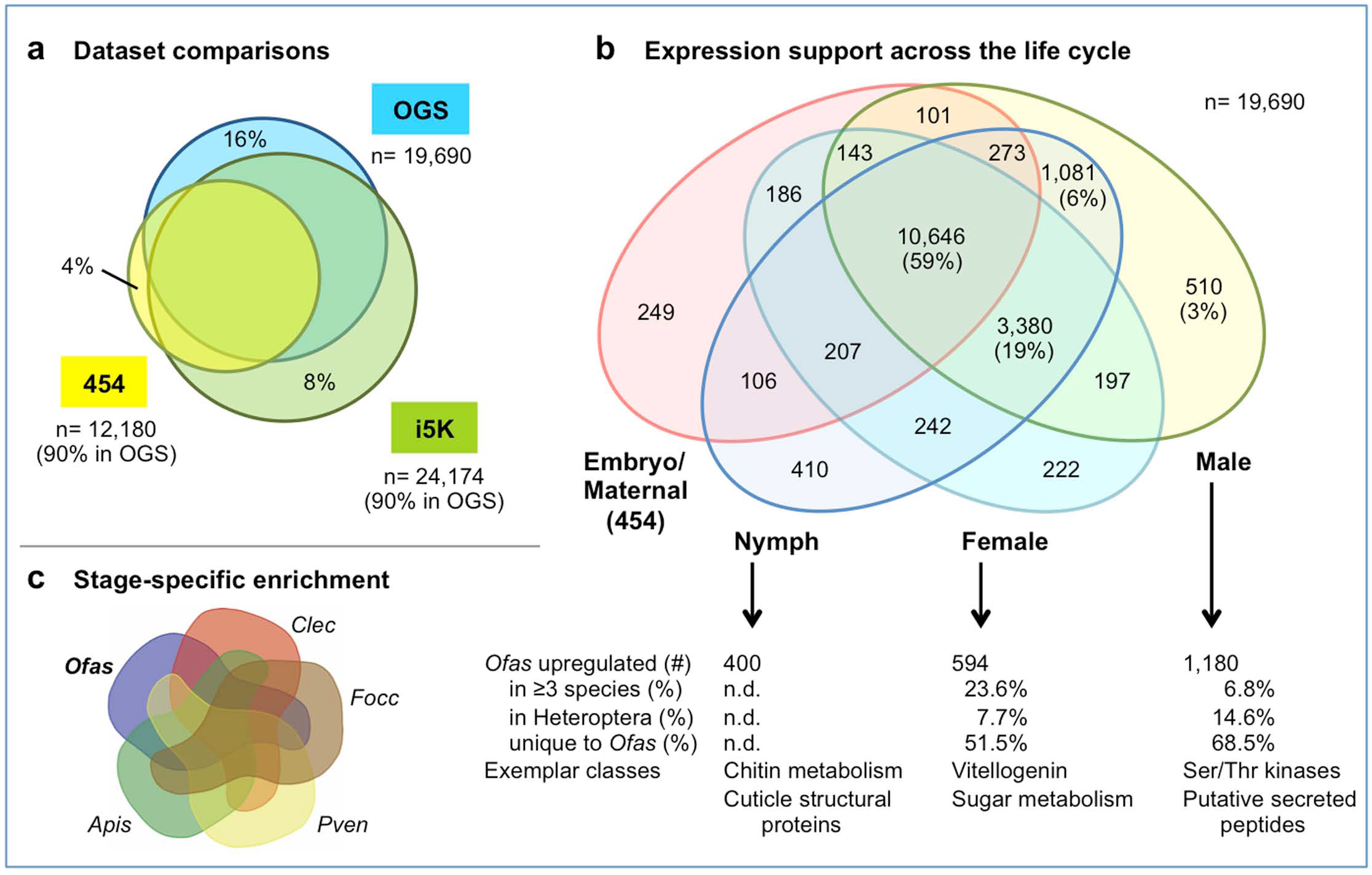
**Comparisons of the official gene set and transcriptomic resources for *Oncopeltus fasciatus*.** **(a)** Area-proportional Venn diagram comparing the OGS v1.1 (“OGS”), a Trinity *de novo* transcriptome from the three post-embryonic RNA-seq samples (“i5K”), and the maternal and embryonic transcriptome from 454 data (“454” [36]). Sample sizes and the fraction of each transcriptome represented in the OGS are indicated (for the 454 dataset, only transcripts with homology identification were considered). The unique fraction of each set is also specified (%). Dataset overlaps were determined by blastn (best hit only, e-value <10^-9^). **(b)** Venn diagram of gene model expression support across four life history samples. Values are numbers of gene models, with percentages also given for the largest subsets. Note that the “Embryo/Maternal” sample derives from 454 pyrosequencing data and therefore has a smaller data volume than the other, Illumina-based samples. **(c)** Summary of sex- and developmental stage-specific RNA-seq comparisons across hemipteroid species: *Acyrthosiphon pisum*, *Apis; Cimex lectularius, Clec; Frankliniella occidentalis, Focc (thysanopteran outgroup); Oncopeltus fasciatus, Ofas; Pachypsylla venusta, Pven*. n.d., not determined. For complete numerical details see Supplemental Note 2.4 Analyses based on OGS v1.1.

The OGS also incorporates many partial and unidentified 454 transcripts, nearly trebling the transcripts with an assigned gene model or homology compared to the original study (from 9% to 26%, by blastn, e<10^-9^). This included 10,130 transcripts that primarily mapped to UTRs and previously lacked recognizable coding sequence, such as for the *Oncopeltus brinker* ortholog, a BMP pathway component ([40], Supplemental Note 5.1.f), and the enzyme-encoding genes *CTP synthase* and *roquin*. At the same time, the transcriptomes provided expression support for the identification of multiple isoforms in the OGS, such as for the germline determinant *nanos* [36]. More generally, most OGS gene models have expression support (91% of 19,690), with 74% expressed broadly in at least three of four samples (Fig. 2b). The inclusion of a fifth dataset from a published adult library [37] provided only a 1% gain in expression support, indicating that with the current study the expression data volume for *Oncopeltus* is quite complete.

RNA-seq studies were further conducted to establish male, female, and nymph specific gene sets (Fig. 2b-c, Supplemental Note 2.4), from which we also infer that the published adult dataset of unspecified sex is probably male. Moreover, most genes with stage-restricted or stage-enriched expression are in our male sample (Fig. 2b-c). For example, gustatory receptor (GR) genes show noticeable restriction to the adult male and published adult (probable male) samples (n= 169 GRs: 40% no expression, 27% only expressed in these two samples), with half of these expressed in both biological replicates (52%). Interestingly, the nymphal sample is enriched for genes encoding structural cuticular proteins (94%, which is >56% more than any other sample). This likely reflects the ongoing molting cycles, with their cyclical upregulation of chitin metabolism and cuticular gene synthesis [41], that are experienced by the different instars and molt cycle stages of individuals pooled in this sample. Lastly, gene sets with sex-specific enrichment across several hemipteroid species substantiate known aspects of male and female reproduction (Fig. 2c: serine-threonine kinases [42] or vitellogenin and other factors associated with oocyte generation, respectively). Some of these enriched genes have unknown functions and could comprise additional, novel factors associated with reproduction in *Oncopeltus*.

### Protein orthology and hemipteran copy number comparisons

To further assay protein-coding gene content, we compared *Oncopeltus* with other arthropods. A phylogeny based on strictly conserved single copy orthologs correctly reconstructs the hemipteran and holometabolan clades’ topologies (Fig. 3a, compare with Fig. 1a), although larger-scale insect relationships remain challenging [3].

**Fig. 3.**
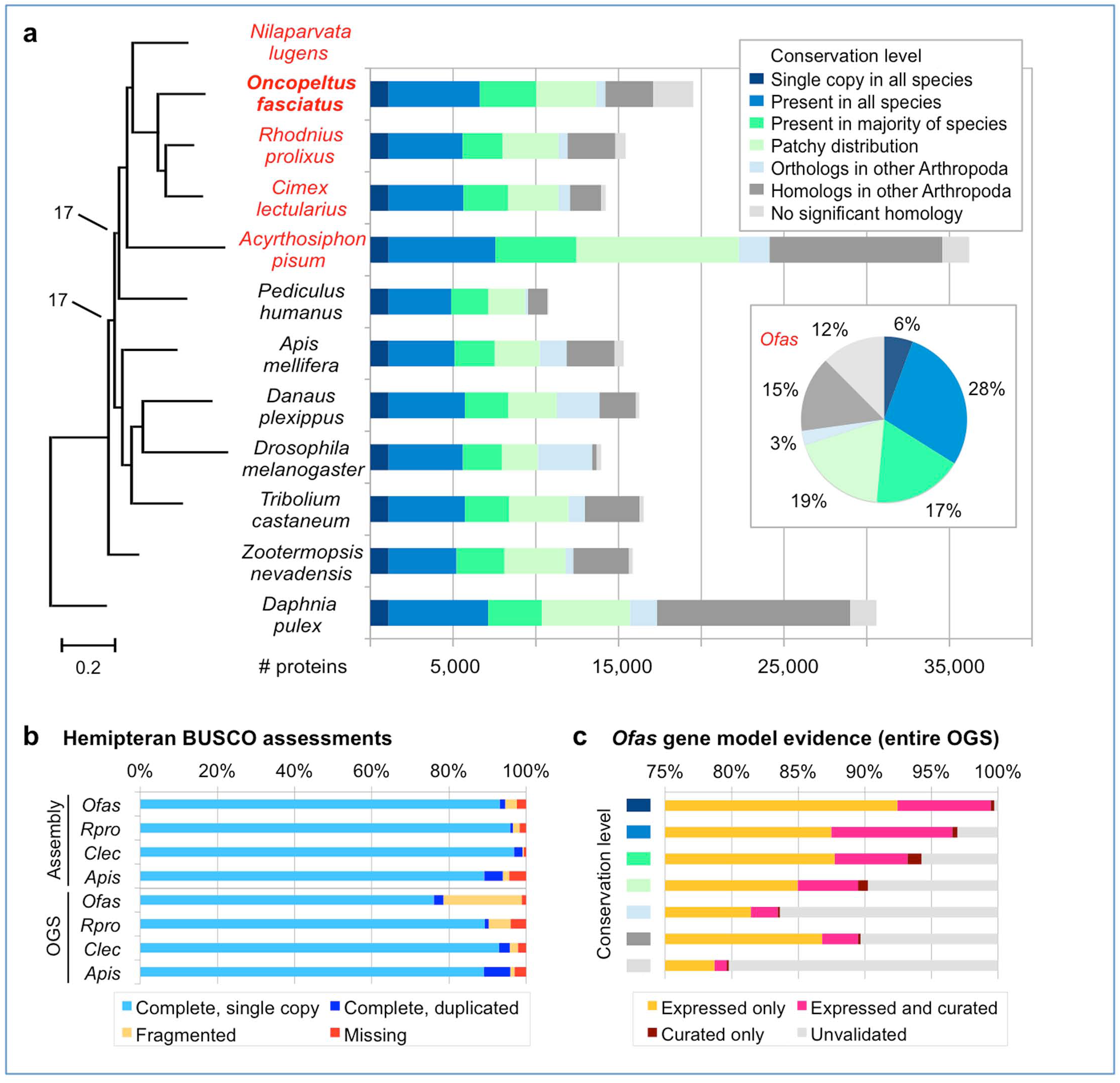
**Orthology comparisons and phylogenetic placement of *Oncopeltus fasciatus* among other Arthropoda.** **(a)** Comparisons of protein-coding genes in 12 arthropod species, with the Hemiptera highlighted in red text. The bar chart shows the number of proteins per conservation level (see legend), based on OrthoDB orthology clustering analyses. To the left is a maximum likelihood phylogeny based on concatenation of 395 single-copy orthologs (all nodes have 100% support unless otherwise noted; branch length unit is substitutions per site). The inset pie chart shows the proportion of proteins per conservation level in *Oncopeltus* (*“Ofas”*). See also Supplemental Note 6.1. **(b)** BUSCO-based analysis of *Oncopeltus* compared to other hemipterans for ortholog presence and copy number in both the assembly and OGS resources, using four-letter species abbreviations (full names in panel a). **(c)** Proportion of *Oncopeltus* proteins that have expression and/or curation validation support per conservation level (same color legend as in (a)). Expression support is based on the life history stage data in Fig. 2b. Analyses based on OGS v1.1.

We then expanded our appraisal to the Benchmarking Universal Single-Copy Orthologs dataset of 1,658 Insecta genes (BUSCO v3, [43]). Virtually all BUSCO genes are present in the *Oncopeltus* OGS (98.9%, Fig. 3b, Supplemental Note 6.1). Although some genes are fragmented, the assembly has a high level of BUSCO completeness (94.6%), independent of annotation prediction limitations that missed some exons from current gene models. Furthermore, BUSCO assessments can elucidate potential consequences of high heterozygosity, which could result in the erroneous inclusion of multiple alleles for a single gene. In fact, the fraction of duplicated BUSCO genes in *Oncopeltus* (1.4%) is low, compared to both the well-assembled bed bug genome (2.2%, [12]) and the pea aphid (4.8%), which is known to have lineage-specific duplications [6, 44]. Thus, by these quality metrics the *Oncopeltus* OGS and assembly are comparable to those of fellow hemipterans, strongly supporting the use of these resources in further comparisons.

We next categorized all proteins by conservation in global, clustering-based orthology analyses (OrthoDB, [1, 45]). As in most species, half of *Oncopeltus* proteins are highly conserved (Fig. 3a). Moreover, 98% of all *Oncopeltus* protein-coding genes have homology, expression, and/or curation support (Fig. 3c). Proteins without homology include species-specific chemoreceptors and antimicrobial peptides (Supplemental Note 5.1.h), as well as potentially novel or partial models. Overall, we estimate that the *Oncopeltus* protein repertoire is comparable to that of other insects in size and conservation. For the Hemiptera, *Oncopeltus* also has fewer missing orthology groups than either the kissing bug or pea aphid (Table S6.1). Indeed the pea aphid is a notable outlier, with its long branch in the phylogeny and for its large protein-coding gene content with low conservation (Fig. 3a). As more hemipteran genomes are sequenced, other species now offer less derived alternatives for phylogenomic comparisons.

Compared to the pea aphid [44], *Oncopeltus* is more conservative in presence and copy number for several signaling pathway components. In contrast to gene absences in the pea aphid, *Oncopeltus* retains orthologs of the EGF pathway component *sprouty*, the BMP receptor *wishful thinking*, and the hormone nuclear receptor *Hr96* (Supplemental Note 5.1.e). Also, whereas multiple copies were reported for the pea aphid, we find a single *Oncopeltus* ortholog for the BMP pathway components *decapentaplegic* and *Medea* and the Wnt pathway intracellular regulator encoded by *shaggy/GSK-3*, albeit with five potential isoforms of the latter (Supplemental Notes 5.1.f, 5.1.j). Duplications of miRNA and piRNA gene silencing factors likewise seem to be restricted to the pea aphid, even compared to other aphid species ([46], Supplemental Note 5.4.a). However, our survey of *Oncopeltus* and other hemimetabolous species reveals evidence for frequent, independent duplications of the Wnt pathway component *armadillo/β-catenin* ([47], Supplemental Note 5.1.j). Curiously, *Oncopeltus* appears to encode fewer histone loci than any other arthropod genome and yet exhibits a similar, but possibly independent, pattern of duplications of histone acetyltransferases to those previously identified in *Cimex* and the pea aphid (Supplemental Note 5.4.c).

On the other hand, we documented several notable *Oncopeltus*-specific duplications. For the BMP transducer *Mad*, we find evidence for three paralogs in *Oncopeltus*, where two occur in tandem and may reflect a particularly recent duplication (Supplemental Note 5.1.f). Similarly, a tandem duplication of *wnt8* appears to be unique to *Oncopeltus* (Supplemental Note 5.1.j). More striking is the identification of six potential paralogs of *cactus*, a member of the Toll/NF-κB signaling pathway for innate immunity, whereas the bed bug and kissing bug each retain only a single copy ([48], Supplemental Note 5.1.g).

Lastly, we explored hemipteran-specific orthology groups against a backdrop of 107 other insect species [1]. What makes a bug a bug in terms of protein-coding genes? Several orthogroups contain potentially novel genes that show no homology outside the Hemiptera and await direct experimental analysis, for which the Hemiptera are particularly amenable (*e.g.*, [49-52, reviewed in 5]). Secondly, there are hemipteran-specific orthogroups of proteins with recognized functional domains and homologs in other insects, but where evolutionary divergence has led to lineage-specific subfamilies. One example is a heteropteran-specific cytochrome P450 (CYP) enzyme (EOG090W0V4B), which in *Oncopeltus* is expressed in all life history stages (Fig. 2b). The expansion of CYP proteins is associated with potential insecticide resistance, as specific P450s can confer resistance to specific chemicals (*e.g.,* [53, 54]; Supplemental Notes 5.3.b, 5.3.c). Hence, the identification of lineage-specific CYP enzymes can suggest potential targets for integrated pest management approaches.

### Transcription factor repertoires and homeobox gene evolution

Having explored the global protein repertoire, we next focused specifically on transcription factors (TFs), which comprise a major class of proteins that has been extensively studied in *Oncopeltus*. This is a class of key regulators of development whose functions can diverge substantially during evolution and for which RNAi-based experimental investigations have been particularly fruitful in the milkweed bug (*e.g.*, [33, 34, 55-57], Supplemental Notes 5.1.a-e).

To systematically evaluate the *Oncopeltus* TF repertoire, we used a pipeline to scan all predicted proteins and assign them to TF families, including orthology assignments where DNA binding motifs could be predicted (see Methods, [58]). We identified 762 putative TFs in *Oncopeltus*, which is similar to other insects for total TF count and for the size of each TF family (Fig. 4a: note that the heatmap also reflects the large, duplicated repertoire in the pea aphid, see also Tables S6.3-S6.5).

**Fig. 4.**
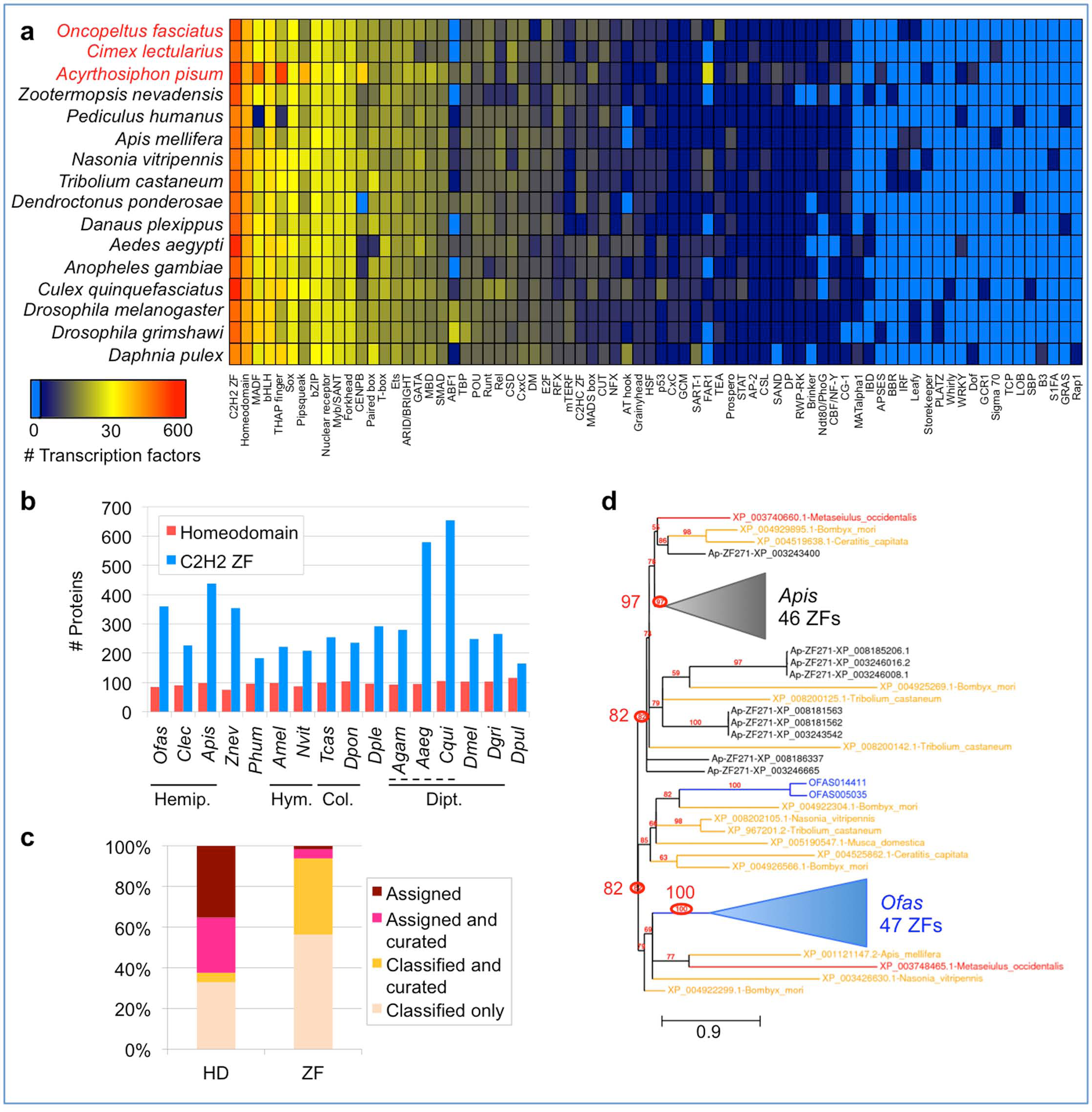
**Distribution of transcription factor families across insect genomes.** **(a)** Heatmap depicting the abundance of 74 transcription factor (TF) families across 16 insect genomes (Hemiptera highlighted in red text), with *Daphnia* as an outgroup, based on the presence of predicted DNA binding domains (see Methods). The color key has a log (base 2) scale (light blue means the TF family is completely absent). Values are in Table S6.3. **(b)** Bar graph showing the number of proteins of each of the two most abundant TF families, homeodomains and C2H2 zinc fingers (ZFs), per species using four-letter abbreviations (full names in panel a). Solid lines demarcate insect orders: Hemiptera (Hemipt.), Hymenoptera (Hym.), Coleoptera (Col.), and Diptera (Dipt.). The dashed line demarcates the dipteran family Culicidae (mosquitoes). **(c)** Proportions of *Oncopeltus* homeodomain (HD) and C2H2 zinc finger proteins with orthology assignment (predicted DNA binding specificity) and/or manual curation. “Classified” refers to automated classification of a protein to a TF family, but without a specific orthology assignment. **(d)** Maximum likelihood phylogeny of representative subsets of the zinc finger 271-like family in *Oncopeltus* (49 proteins, blue text) and the pea aphid (55 proteins, black text), with chelicerate (red text) and holometabolan (yellow text) outgroups (16 proteins, 7 species), based on the *Oncopeltus* OGS and GenBank protein accessions. Gaps were removed during sequence alignment curation; all nodes have ≥50% support; branch length unit is substitutions per site [158]. Key nodes are circled for the clades containing all aphid or all *Oncopeltus* proteins (82% support each), and for each ‘core’ clade comprised exclusively of proteins from each species (97% and 100%, respectively; triangles shown to scale for branch length and number of clade members). Branch length unit is substitutions per site. Analyses based on OGS v1.1.

We were able to infer DNA binding motifs for 25% (n=189) of *Oncopeltus* TFs, mostly based on data from *Drosophila melanogaster* (121 TFs) but also from distantly related taxa such as mammals (56 TFs). Such high conservation is further reflected in explicit orthology assignments for most proteins within several large TF families, including the homeodomain (53 of 85, 62%), basic helix-loop-helix (bHLH, 35 of 45, 78%), and forkhead box (16 of 17, 94%) families. In contrast, most C2H2 zinc finger proteins lack orthology assignment (only 22 of 360, 6%). Across species, the homeodomain and C2H2 zinc finger proteins are the two largest TF superfamilies (Fig. 4a). Given their very different rates of orthology assignment, we probed further into their pipeline predictions and the patterns of evolutionary diversification.

The number of homeodomain proteins identified by the pipeline displays a narrow normal distribution across species (Fig. 4b, mean ± standard deviation: 97 ± 9), consistent with a highly conserved, slowly evolving protein family. Supporting this, many *Oncopeltus* homeodomain proteins that were manually curated also received a clear orthology assignment (Fig. 4c: pink), with only four exceptions (Fig. 4c: yellow). Only one case suggests a limitation of a pipeline that is not specifically tuned to hemipteran proteins (Goosecoid). Manual curation of partial or split models identified a further 11 genes encoding homeodomains, bringing the actual tally in *Oncopeltus* to 96. Overall, we find the TF pipeline results to be a robust and reasonably comprehensive representation of these gene classes in *Oncopeltus*.

These analyses also uncovered a correction to the published *Oncopeltus* literature for the developmental patterning proteins encoded by the paralogs *engrailed* and *invected*. These genes arose from an ancient tandem duplication prior to the hexapod radiation. Their tail-to-tail orientation enables ongoing gene conversion [59], making orthology discrimination particularly challenging. For *Oncopeltus*, we find that the genes also occur in a tail-to-tail orientation and that *invected* retains a diagnostic alternative exon [59]. These new data reveal that the purported *Oncopeltus engrailed* ortholog in previous developmental studies (*e.g.*, [55, 60-63]) is in fact *invected* (Supplemental Note 5.1.a).

### Independent expansions of C2H2 zinc fingers within the Hemiptera

Unlike homeodomain proteins, C2H2 zinc finger (C2H2-ZF) repertoires are prominent for their large family size and variability throughout the animal kingdom [64], and this is further supported by our current analysis in insects. With >350 C2H2-ZFs, *Oncopeltus*, the pea aphid, termite, and some mosquito species have 1.5× more members than the insect median (Fig. 4b). This is nearly half of all *Oncopeltus* TFs. While the expansion in mosquitoes could have a single origin within the Culicinae, the distribution in the Hemiptera, where *Cimex* has only 227 C2H2-ZFs, suggests that independent expansions occurred in *Oncopeltus* and the pea aphid. Prior to the sequencing of other hemipteran genomes, the pea aphid’s large C2H2-ZF repertoire was attributed to the expansion of a novel subfamily, APEZ, also referred to as zinc finger 271-like [44].

In fact, manual curation in *Oncopeltus* confirms the presence of a subfamily with similar characteristics to APEZ (Fig. 4c: yellow fraction). In *Oncopeltus* we find >115 proteins of the ZF271 class that are characterized by numerous tandem repeats of the C2H2-ZF domain and its penta-peptide linker, with 3-45 repeats per protein.

Intriguingly, we find evidence for ongoing evolutionary diversification of this subfamily. A number of *Oncopeltus* ZF271-like genes occur in tandem clusters of 4-8 genes – suggesting recent duplication events. Yet, clustered genes differ in gene structure (number and size of exons), and we identified a number of probable ZF271-like pseudogenes whose open reading frames have become disrupted – consistent with high turnover. *Oncopeltus* ZF271-like proteins also differ in the sequence and length of the zinc finger domains among themselves and compared to aphid proteins (WebLogo analysis, [65]), similar to zinc finger array shuffling seen in humans [66]. Furthermore, whole-protein phylogenetic analysis supports independent, rapid expansions in the pea aphid and *Oncopeltus* (Fig. 4d).

Clustered zinc finger gene expansion has long been recognized in mammals, with evidence for strong positive selection to increase both the number and diversity of zinc finger domains per protein as well as the total number of proteins [67]. This was initially found to reflect an arms-race dynamic of co-evolution between selfish transposable elements and the C2H2-ZF proteins that would repress them [68]. In vertebrates, these C2H2-ZF proteins bind to the promoters of transposable elements via their zinc finger arrays and use their Krüppel-associated box (KRAB) domain to bind the chromatin-remodeling co-repressor KAP-1, which in turn recruits methyltransferases and deacetylases that silence the targeted promoter [69].

Insects do not have a direct ortholog of vertebrate KAP-1 (Supplemental Note 5.4.d), and neither the aphid nor *Oncopeltus* ZF271-like subfamilies possess a KRAB domain or any other domain besides the zinc finger arrays. However, close molecular outgroups to this ZF271-like subfamily include the developmental repressor Krüppel [70] and the insulator protein CTCF [71] (data not shown). Like these outgroups, the *Oncopeltus* ZF271-like genes are strongly expressed: 98% have expression support, with 86% expressed in at least three different life history stages (Fig. 2b). Thus, the insect ZF271-like proteins may also play prominent roles in repressive DNA binding. Indeed, we find evidence for a functional methylation system in *Oncopeltus* (Supplemental Note 5.4.c), like the pea aphid, which would provide a means of gene silencing by chromatin remodeling, albeit via mediators other than KAP-1.

However, an arms race model need not be the selective pressure that favors insect ZF271-like family expansions. Recent analyses in vertebrates identified sophisticated, additional regulatory potential by C2H2-ZF proteins, building upon original transposable element binding for new, lineage-specific and even positive gene regulation roles [66, 72, 73]. Moreover, although *Cimex* has half as many long terminal repeat (LTR) repetitive elements as *Oncopeltus* and the pea aphid, overall we do not find a correlation between relative or absolute repetitive content and ZF271-like family expansion within the Hemiptera (see next section).

### Proportional repeat content across hemipterans

With the aim of reducing assembly fragmentation and to obtain a better picture of repeat content, we performed low coverage, long read PacBio sequencing in *Oncopeltus* (Supplemental Note 2.3). Using PacBio reads in a gap-filling assay on the Illumina assembly raised the total detected repetitive content from 25% to 32%, while repeat estimations based on simultaneous assessment of Illumina and PacBio reads nearly doubled this value to 58%. As expected, the capacity to identify repeats is strongly dependent on assembly quality and sequencing technology, with the *Oncopeltus* repetitive content underrepresented in the current (Illumina-only) assembly. Furthermore, as increasing genome size compounds the challenge of assembling repeats, the repeat content of the current assembly is lower than in species with smaller genome sizes (Fig. 5a, with the sole exception of the honey bee), and we therefore used our gap-filled dataset as a more accurate basis for further comparisons.

**Fig. 5.**
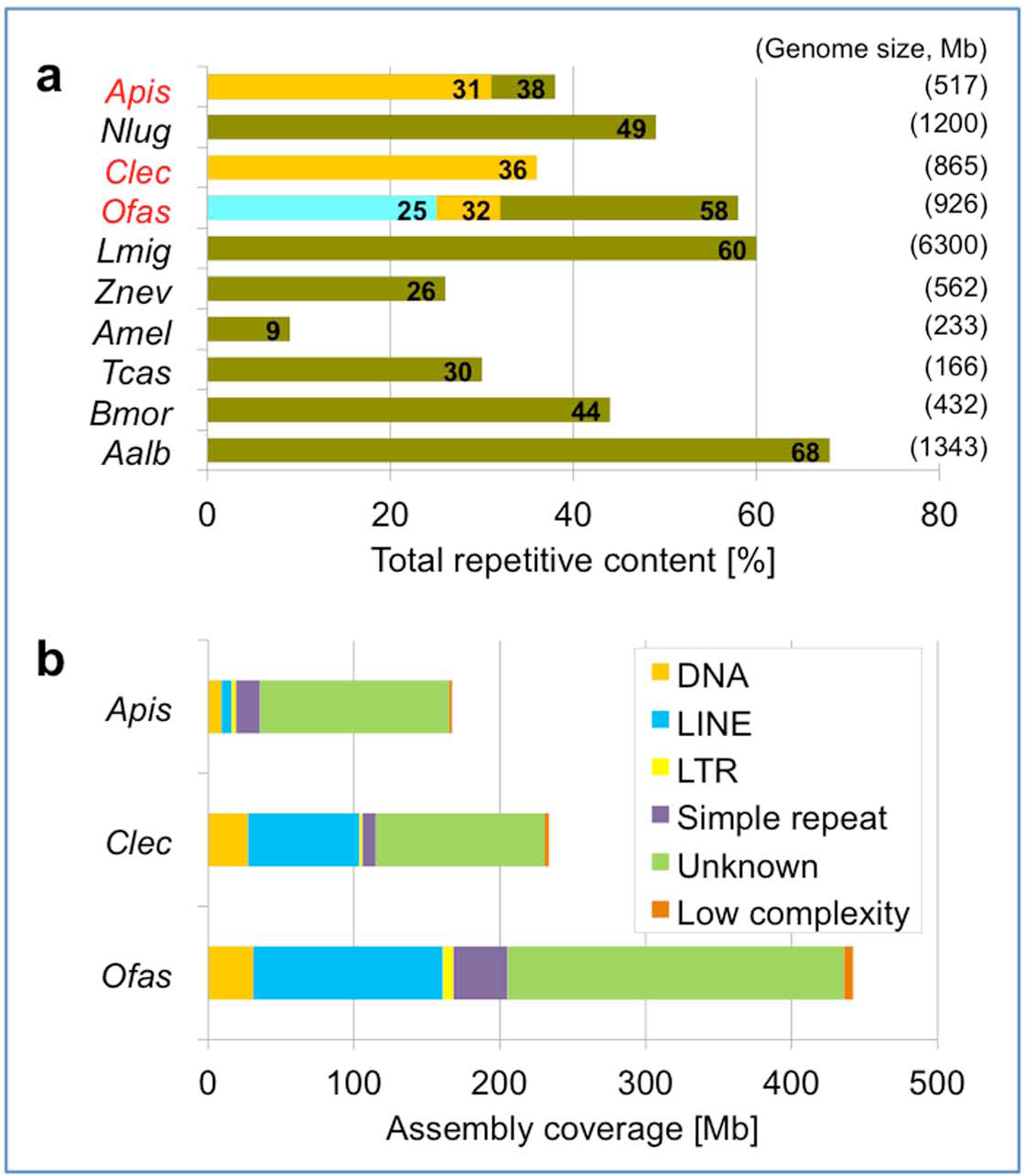
**Comparison of repeat content estimations.** **(a)** Comparison of total repetitive content among insect genomes. The three values for *Oncopeltus* are shown (in ascending order: original Illumina assembly, gap-filled assembly, Illumina-PacBio hybrid estimate). Values for the three hemipterans labeled in red text are from RepeatModeler (gold bars for the pea aphid and bed bug; blue and gold bars for *Oncopeltus*). All other values are from the respective genome papers, including a second value corresponding to the published repeat content for the first version of the aphid genome [6, 10, 106, 159-164]. Species abbreviations as in Fig. 4, and additionally: *Nlug, Nilaparvata lugens; Lmig, Locusta migratoria; Bmor, Bombyx mori; Aalb, Aedes albopictus*. **(b)** Comparison of repetitive element categories between three hemipteran genomes, based on results from RepeatModeler. Here we present assembly coverage as actual sequence length (Mb) to emphasize the greater repeat content in *Oncopeltus* (based on the gap-filled assembly, see also Supplemental Note 2.3).

To support direct comparisons among hemipterans, we also performed our RepeatModeler analysis on the bed bug and pea aphid assemblies. Repeats comprised 36% and 31% of the respective assemblies, similar to the gap-filled value of 32% in *Oncopeltus*. Nevertheless, given the smaller sizes of these species’ assemblies – 651 Mb in the bed bug and 542 Mb in the pea aphid – the absolute repeat content is much higher in *Oncopeltus* (Fig. 5b). Excluding unknown repeats, the most abundant transposable elements in *Oncopeltus* are LINE retrotransposons, covering 10% of the assembly (Table S2.5). This is also the case in the bed bug (12%), while in the pea aphid DNA transposons with terminal inverted repeats (TIRs) are the most abundant (2% of the assembly identified here, and 4% reported from manual curation in the pea aphid genome paper, [6]). Across species, the remaining repeat categories appear to grow proportionally with assembly size, except for simple repeats, which were the category with the largest relative increase in size after gap-filling in *Oncopeltus* (Supplemental Note 2.3). However, given the mix of data types (Illumina [12] and Sanger [6]), these patterns should be treated as hypotheses for future testing.

### Lineage and genome size-related trends in insect gene structure

Both our manual curation work and BUSCO analyses highlighted the fact that *Oncopeltus* genes are often comprised of many, small exons. We thus undertook a comparative analysis to determine whether this is a general feature to be considered for structural annotation of hemipteran genomes. We find that both lineage and genome size can serve as predictors of gene structure.

Firstly, we created a high quality dataset of 30 functionally diverse, large genes whose manual curation could reasonably ensure complete gene models across seven species from four insect orders (Fig. 6a, Supplemental Note 6.3). Most species encode the same total number of amino acids for these conserved proteins, with the thrips *Frankliniella occidentalis* and the fruit fly being notable exceptions with larger proteins (Fig. 6a: blue plot line). However, the means of encoding this information differs between lineages, with hemipteroid orthologs comprised of twice as many exons as their holometabolous counterparts (Fig. 6a: orange plot line). Thus, there is an inverse correlation between exon number and exon size (Fig. 6a: orange vs. red plot lines). This analysis corroborates and extends previous probabilistic estimates of intron density, where the pea aphid as a sole hemipteran representative had the highest intron density of ten insect species [74].

**Figure 6.**
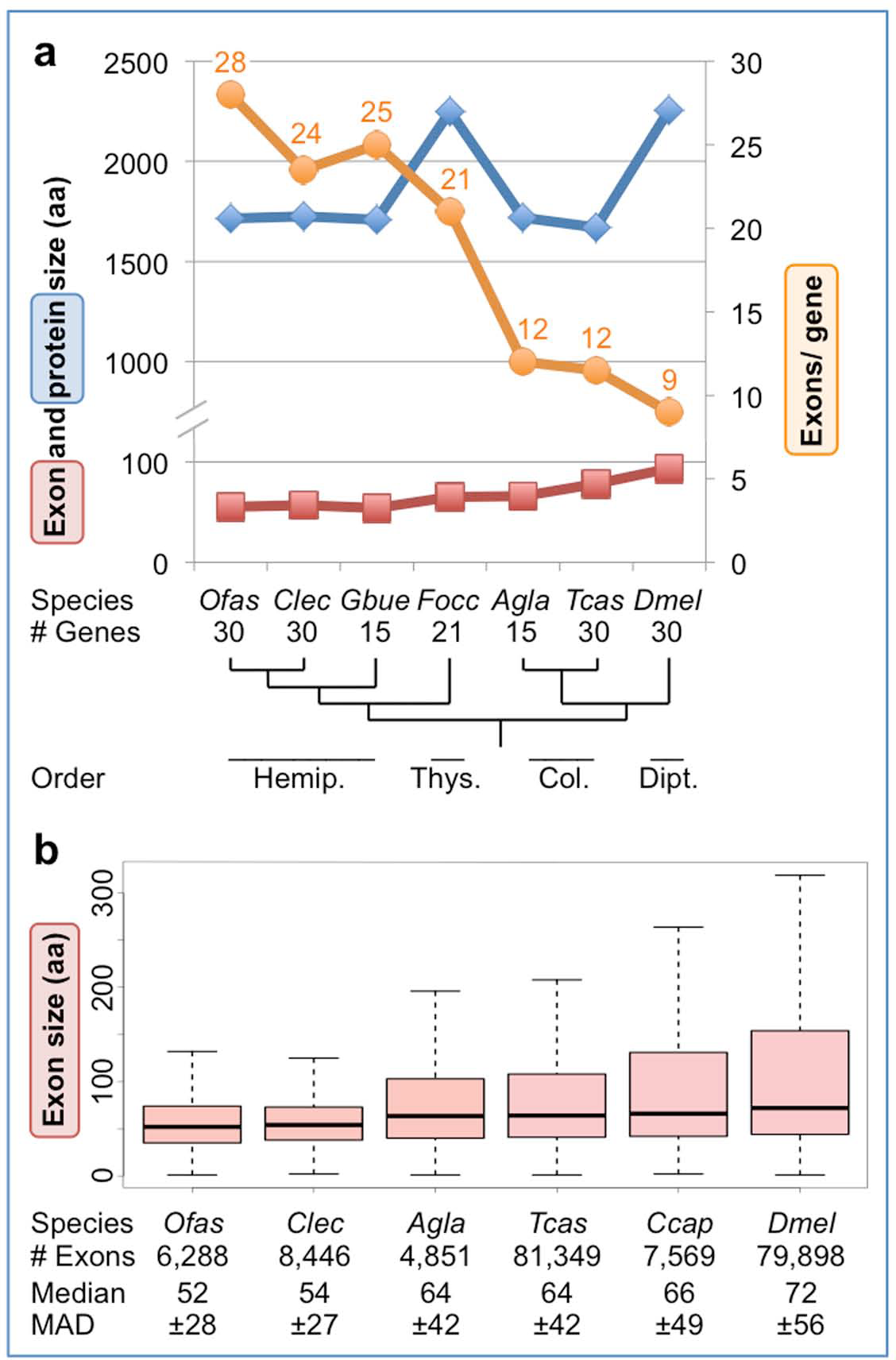
**Trends in gene structure show hemipteroid-specific tendencies.** **(a)** Median values per species for protein size, exon size, and exon number for a curated set of highly conserved genes encoding large proteins of diverse functional classes (see also Supplemental Note 6.3). Sample sizes are indicated, with 11 genes for which orthologs were evaluated in all species. Where it was not possible to analyze all 30 genes for a given species, equal sampling was done across the range of protein sizes of the complete dataset, based on the *Cimex* ortholog sizes (1:1:1 sampling from big:medium:small subcategories of 10 genes each). **(b)** Box plot representations of coding sequence exon size (aa) for two species from each of three insect orders, based on datasets of unique coding sequence exons (one isoform per gene) and excluding terminal exons <10 aa (as most of those exons may rather be UTRs or a small placeholder N-terminal exon based on automated Maker model predictions). Only manually curated gene models were considered for the i5K species, including *Oncopeltus*; the entire OGS was used for *Tribolium* and *Drosophila*. For clarity, outliers are omitted; whiskers represent 1.5× the value of the Q3 (upper) or Q2 (lower) quartile range. MAD, median absolute deviation. Species are represented by their four-letter abbreviations, with their ordinal relationships given below the phylogeny in panel (a): Hemip., Hemiptera; Thys., Thysanoptera; Col., Coleoptera; Dipt., Diptera. Species abbreviations as in Figs. 2,4 and additionally: *Gbue, Gerris buenoi* [165]; *Agla, Anoplophora glabripennis* [30]; *Ccap, Ceratitis capitata* [166].

To test these trends, we next expanded our analysis to all manually curated exons in two species from each of three orders (Hemiptera, Coleoptera, Diptera). Here, we expect that curated exon sizes are accurate, without the need to assume that entire gene models are complete. This large dataset corroborates our original findings, with bugs having small exons while both the median and Q3 quartile reflect larger exon sizes in beetles and flies (Fig. 6b). Notably, the median and median absolute deviation are highly similar between species pairs within the Hemiptera and Coleoptera. Meanwhile, the exon metrics within the Diptera support large protein sizes as a drosophilid-specific, rather than dipteran-wide, feature.

Does the high exon count in the Hemiptera reflect an ancient, conserved increase at the base of this lineage, or ongoing remodeling of gene structure with high turnover? To assess the exact nature of evolutionary changes, we annotated intron positions within multiple sequence alignments of selected proteins and plotted gains and losses onto the phylogeny, providing a total sample of 165 evolutionary changes at 148 discrete splice sites (Fig. 7, see also Supplemental Note 6.3 for gene selection and method). These data reveal several major correlates with intron gain or loss. The bases of both the hemipteroid and hemipteran radiations show the largest gains, while most losses occur in the dipteran lineage (Fig. 7: orange and purple shading, respectively). Furthermore, we find progressive gains across hemipteroid nodes, and it is only in this lineage that we additionally find species-specific splice changes for the highly conserved *epimerase* gene (Fig. 7: orange outline). Thus, we find evidence for both ancient intron gain and ongoing gene structure remodeling in this lineage.

**Fig 7.**
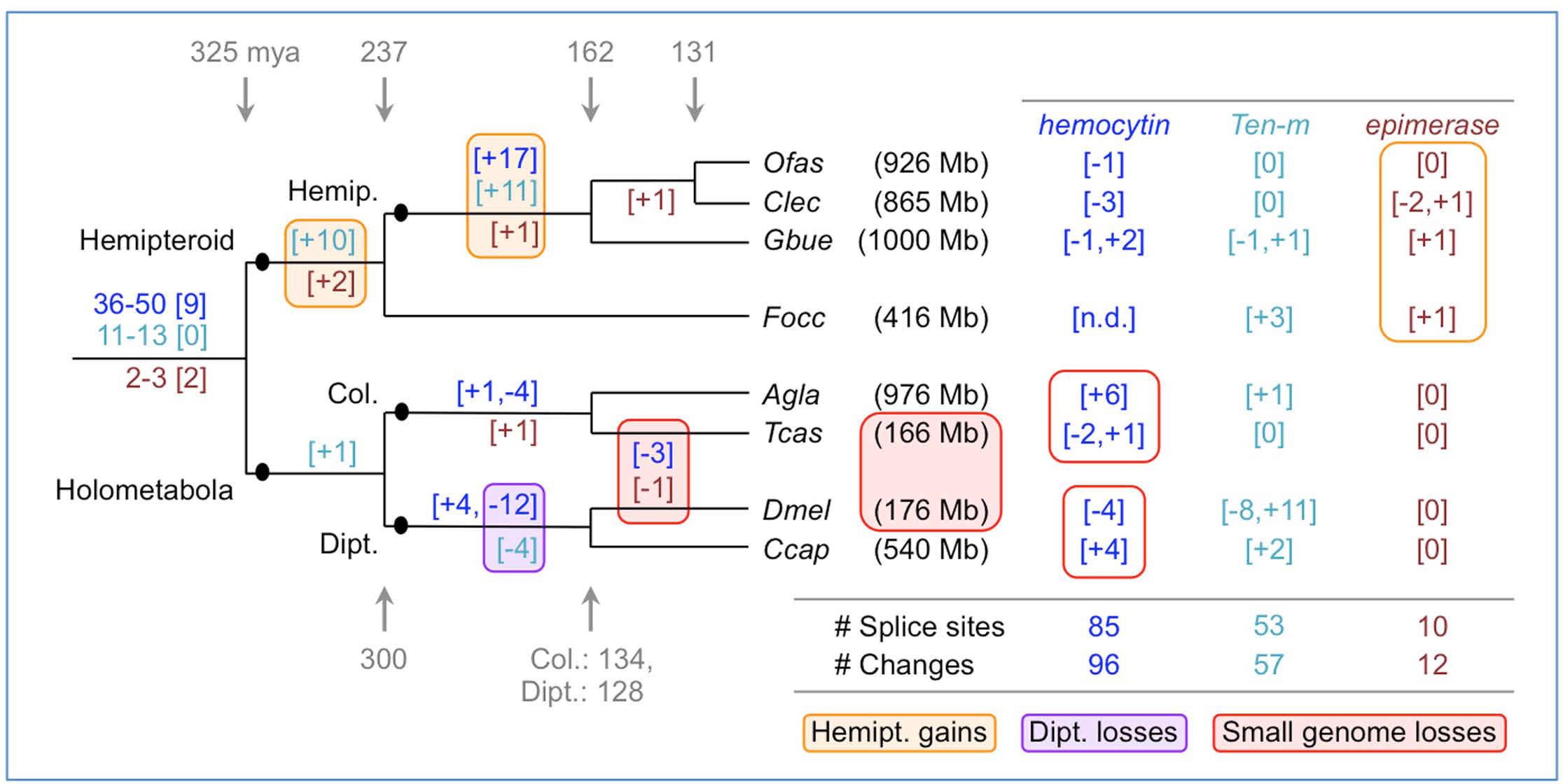
**Splice site evolution correlates with both lineage and genome size.** Splice site changes are shown for *hemocytin* (blue text), *Tenascin major* (*Ten-m*, turquoise text), and *UDP-galactose 4≠-epimerase* (brown text), mapped onto a species tree of eight insects. Patterns of splice site evolution were inferred based on the most parsimonious changes that could generate the given pattern within a protein sequence alignment of all orthologs (see also Supplemental Note 6.3 for methodology and data sources). If inferred gains or losses were equally parsimonious, we remained agnostic and present a range for the ancestral number of splice sites present at the base of the tree, where the bracketed number indicates how many ancestral positions are still retained in all species. Along each lineage, subsequent changes are indicated in brackets, with the sign indicating gains (+) or losses (-). Values shown to the right are species-specific changes. The values shown between the *D. melanogaster* and *T. castaneum* lineages denote changes that have occurred independently in both species. Colored boxes highlight the largest sources of change, as indicated in the legend. Species are represented by their four-letter abbreviations (as in Fig. 6), and estimated genome sizes are indicated parenthetically (measured size: [12, 30, 163, 166, 167]; draft assembly size: GenBank Genome IDs 14741 and 17730). Divergence times are shown in gray and given in millions of years [3]. Abbreviations as in Figs. 4,6, and also: Col., Coleoptera; Dipt., Diptera; Hemip., Hemiptera; Hemipt., hemipteroid assemblage (including *F. occidentalis*); n.d., no data.

Surprisingly, both *hemocytin* and *epimerase* – our exemplar genes with many (up to 74) and few exons (3-8 per species), respectively – show independent losses of the same splice sites in *Drosophila* and *Tribolium*. One feature these species share is a genome size 2.4-6.0× smaller than in the other species examined here (Fig. 7: red shading). Pairwise comparisons within orders also support this trend, as the beetle and fly species with larger genomes exhibit species-specific gains compared to intron loss in their sister taxa (Fig. 7: red outlines). Thus, genome size seems to positively correlate with intron number. However, lineage is a stronger predictor of gene structure: the coleopteran and dipteran species pairs have highly similar exon size metrics despite differences in genome size (Fig. 6b). A global computational analysis over longer evolutionary distances also supports a link between genome size and intron number in arthropods, although chelicerates and insects may experience different rates of evolutionary change in these features [75]. It will be interesting to see if the correlation with genome size is borne out in other invertebrate taxa.

The selective pressures and mechanisms of intron gain in the Hemiptera will be a challenge to uncover. While median exon size (Fig. 6b) could reflect species-specific nucleosome sizes [76, 77], this does not explain why only the Hemiptera seldom exceed this (Fig. 6b: Q3 quartile). Given gaps in draft genome assemblies, we remain cautious about interpreting (large) intron lengths but note that many hemipteran introns are too small to have harbored a functional transposase gene (*e.g.,* median intron size of 429 bp, n=69 introns in *hemocytin* in *Cimex*). Such small introns could be consistent with proliferation of non-autonomous short interspersed nuclear elements (SINEs). However, characterization of such highly divergent non-coding elements would require curated SINE libraries for insects, comparable to those generated for vertebrates and plants [76, 77]. Meanwhile, it appears that hemipteran open reading frames ≥160 bp are generally prevented by numerous in-frame stop codons just after the donor splice site. Most stop codons are encoded by the triplet TAA in both *Oncopeltus* and *Cimex* (data not shown), although these species’ genomes are not particularly AT-rich (Table 1).

Even if introns are small, having gene loci comprised of numerous introns and exons adds to the cost of gene expression in terms of both transcription duration and mRNA processing. One could argue that a gene like *hemocytin*, which encodes a clotting agent, would require rapid expression in the case of wounding – a common occurrence in adult *Cimex* females due to the traumatic insemination method of reproduction [12]. Thus, as our molecular understanding of comparative insect and particularly hemipteran biology deepens, we will need to increasingly consider how life history traits are manifest in genomic signatures at the structural level (*e.g.*, Figs. 5-7), as well as in terms of protein repertoires (Figs. 3-4).

### Expansion after a novel lateral gene transfer (LGT) event in phytophagous bugs

In addition to the need for cuticle repair, traumatic insemination may be responsible for the numerous LGT events predicted in the bed bug [12]. In contrast, the same pipeline analyses [78] followed by manual curation predicted very few LGTs in *Oncopeltus*, which lacks this unusual mating behavior. Here, we have identified 11 strong LGT candidates, and we confirmed the incorporation of bacterial DNA into the milkweed bug genome for all five candidates chosen for empirical testing (Table S2.4). Curiously, we find several LGTs potentially involved in bacterial or plant cell wall metabolism that were acquired from different bacterial sources at different times during hemipteran lineage evolution, including two distinct LGTs that are unique to *Oncopeltus* and implicated in the synthesis of peptidoglycan, a bacterial cell wall constituent (Supplemental Note 2.2).

Conversely, two other validated LGT candidates are implicated in cell wall degradation. We find two strongly expressed, paralogous copies in *Oncopeltus* of a probable bacterial-origin gene encoding an endo-1,4-beta-mannosidase enzyme (MAN4, EC 3.2.1.78). Inspection of genome assemblies and protein accessions reveals that this LGT event occurred after the infraorder Pentatomomorpha, including the stink bug *Halyomorpha halys*, diverged from other hemipterans, including the bed bug (Fig. 8a). Independent duplications then led to the two copies in *Oncopeltus* and an astonishing nine tandem copies in *Halyomorpha* (Figs. 8b, S2.6). Since the original LGT event, the *mannosidase* genes have gained introns that are unique to each species and to subsets of paralogs (Fig. 8c). Thus, the “domestication” [79] of *mannosidase* homologs as multi-exonic genes further illustrates the hemipteran penchant for intron introduction and maintenance of small exons. The retention and subsequent expansion of these genes implies their positive selection, consistent with the phytophagous diet of these species. It is tempting to speculate that copy number proliferation in the stink bug correlates with the breadth of its diet, as this agricultural pest feeds on a number of different tissues in a range of host plants [80].

**Fig. 8.**
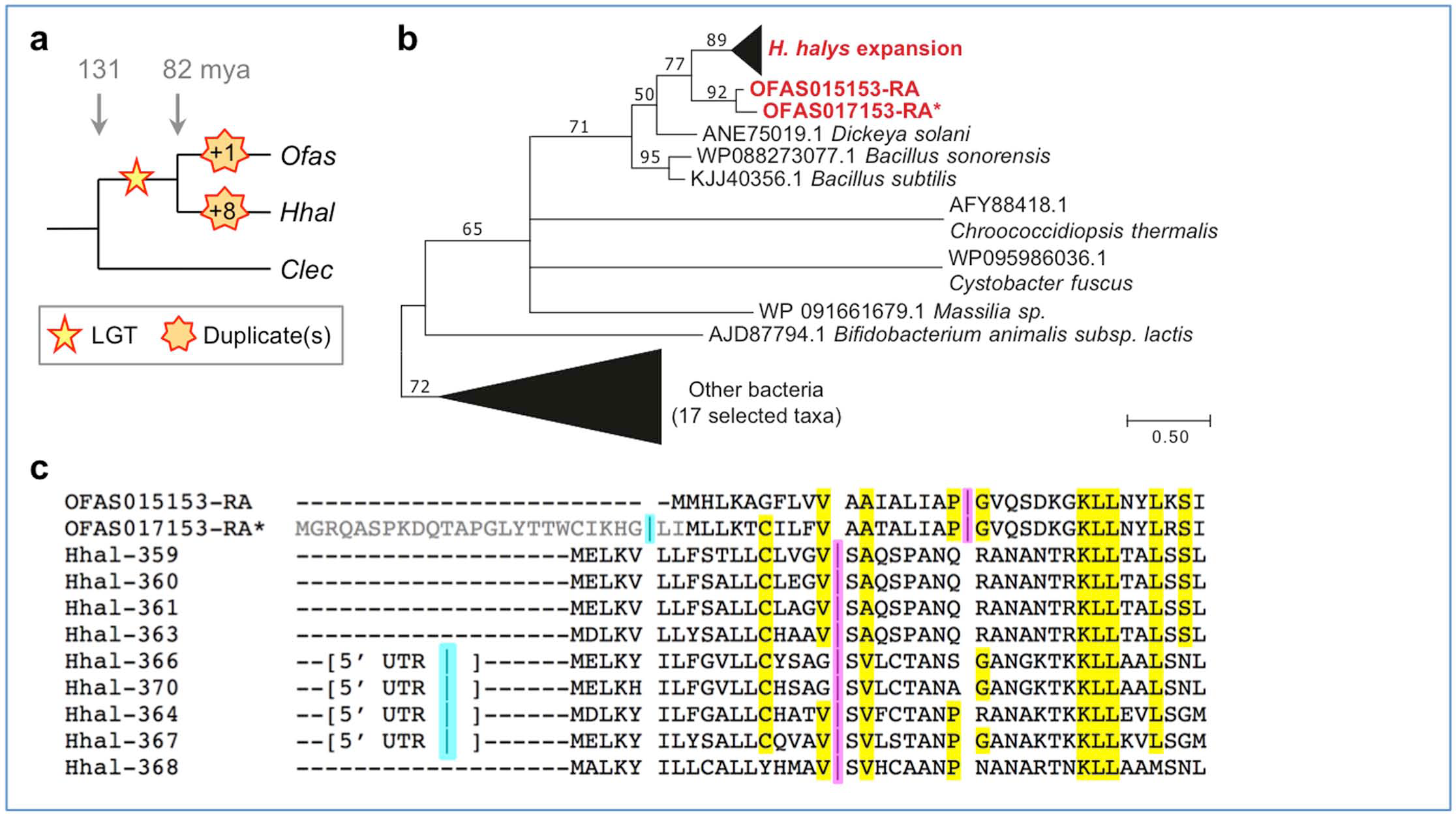
**Lateral gene transfer introduction and subsequent evolution within the Hemiptera for mannosidase-encoding genes.** **(a)** Species tree summary of evolutionary events. Stars represent the original LGT introduction and subsequent copy number gains (see legend). **(b)** Maximum likelihood phylogeny of mannosidase proteins, including bacterial sequences identified among the best GenBank blastp hits for *Oncopeltus* and *Halyomorpha* (accession numbers as indicated, and for “Other bacteria” are: ACB22214.1, AEE17431.1, AEI12929.1, AEO43249.1, AFN74531.1, CDM56239.1, CUA67033.1, KOE98396.1, KPI24888.1, OAN41395.1, ODP26899.1, ODS11151.1, OON18663.1, PBD05534.1, SIR54690.1, WP096035621.1, YP001327394.1). All nodes have ≥50% support from 500 bootstrap replicates [168]. Triangles are shown to scale for branch length and number of clade members; branch length unit is substitutions per site. See also Fig. S2.6. **(c)** Manually curated protein sequence alignment for the N-terminal region only. Splice sites (“|” symbol) are shown, where one position is ancestral and present in all paralogs of a given species (magenta) and one position occurs in a subset of paralogs and is presumed to be younger (cyan, within the 5≠ UTR in *Halyomorpha*). Residues highlighted in yellow are conserved between the two hemipteran species. The *Oncopeltus* paralog represented in the OGS as OFAS017153-RA is marked with an asterisk to indicate that this version of the gene model is incomplete and lacks the initial exon (gray text in the alignment). For clarity, only the final three digits of the *Halyomorpha* GenBank accessions are shown (full accessions: XP_014289XXX).

### Cuticle development, structure, and warning pigmentation

Body cuticle is another trait associated with feeding ecology, particularly for pigmentation. Furthermore, the milkweed bug has been a powerful model for endocrine studies of hemimetabolous molting and metamorphosis since the 1960’s [22, 81-84]. Therefore, we next focused on the presence and function of genes involved in cuticle development and structure.

Molting is triggered by the release of ecdysteroids, steroid hormones that are synthesized from cholesterol by cytochrome P450 enzymes of the Halloween family [85], and we were able to identify these in the *Oncopeltus* genome (Supplemental Notes 5.2.b, 5.3.b). From the ecdysone response cascade defined in *Drosophila* [86], we identified *Oncopeltus* orthologs of both early and late-acting factors, including ecdysteroid hormones and their receptors. It will be interesting to see if the same regulatory relationships are conserved in the context of hemimetabolous molting in *Oncopeltus*. For example, *E75A* is required for reactivation of ecdysteroid production during the molt cycle in *Drosophila* larvae [87] and likely operates similarly in *Oncopeltus*, since *Of-E75A* RNAi prevents fourth-instar nymphs from molting to the fifth instar (H. Kelstrup and L. Riddiford, unpublished data).

In hemipterans, activation of juvenile hormone (JH) signaling at molts determines whether the insect progresses to another nymphal instar or, if lacking, becomes an adult [50]. We were able to identify many components of the JH signal transduction pathway in the *Oncopeltus* genome, including orthologs of *Methoprene-tolerant* (*Met*), the JH receptor [50, 88], and the JH-response gene *Kr-h1* [50, 89, 90]. JH acts to determine cuticle identity through regulation of the *broad* gene in a wide variety of insects, where different isoforms direct specific aspects of metamorphosis in *Drosophila* [91, 92]. In *Oncopeltus*, *broad* expression directs progression through the nymphal stages [93], but the effect of each isoform was unknown. We identified three isoforms in *Oncopeltus* – *Z2*, *Z3*, and *Z4* – and performed isoform-specific RNAi. In contrast to *Drosophila*, Broad isoform functions appear to be more redundant in *Oncopeltus*, as knockdown of isoforms *Z2* and *Z3* has similar effects on survival to adulthood as well as adult wing size and morphology (Fig. 9).

**Fig. 9.**
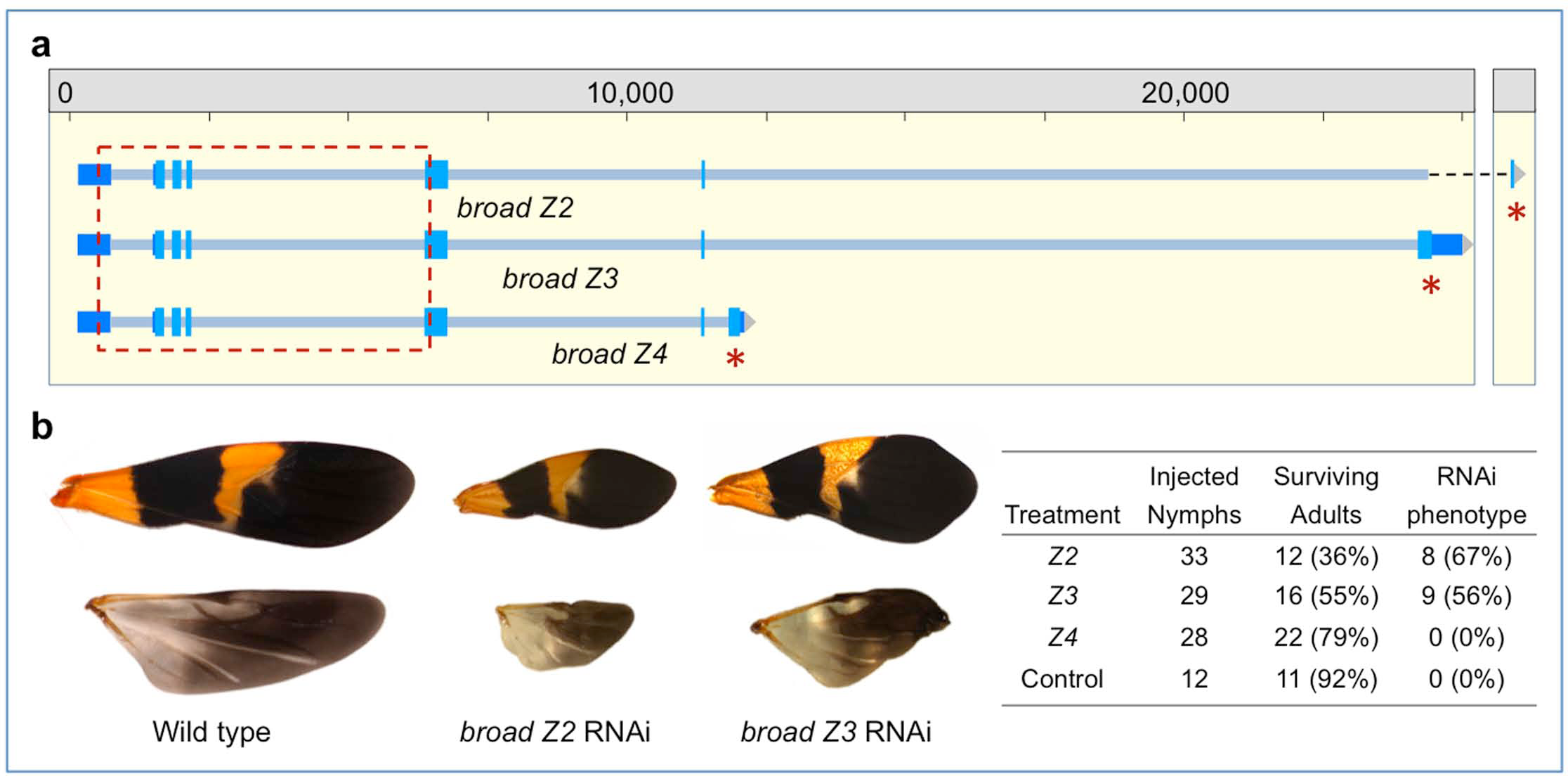
**Isoform-specific RNAi based on new genome annotations affects the molting and cuticle identity gene *broad*.** **(a)** Genomic organization of the cuticle identity gene *broad*. The regions used as template to generate isoform-specific dsRNA are indicated (red asterisks: the final, unique exons of each isoform). Previous RNAi studies targeted sequence within exons 1-5 that is shared among all isoforms (dashed red box, [93]). **(b)** Knock down of the *Oncopeltus Z2* or *Z3 broad* isoforms at the onset of the penultimate instar resulted in altered nymphal survival and morphogenesis that was reflected in the size and proportion of the fore and hind wings at the adult stage (upper and lower images, respectively, shown to the same scale for all wings). We did not detect any effect on the wing phenotype when targeting the *Z4*-specific exon, demonstrating the specificity of the zinc finger coding region targeted by RNAi. Experimental statistics are provided in the figure inset, including for the buffer-injected negative control.

Regulators such as Broad initiate the transcription of a large battery of genes that encode the structural components of the cuticle needed at each molt, consistent with our expression analyses (Fig. 2b-c, discussed above). We identified 173 genes encoding putative cuticle structural proteins in *Oncopeltus* (Supplemental Note 5.2.c). Similar to other insects, the CPR family, with the RR-1 (soft cuticle), RR-2 (hard cuticle), and unclassifiable types, constituted the largest cuticle protein family. While several protein families are similar in size to those of other insects (CPAP1, CPAP3, and TWDL: Table S5.12), we found a slight expansion in the *Oncopeltus* CPF family (Fig. S5.14). For cuticle production, similar to the bed bug and the Asian longhorned beetle [12, 30], we identified a single *chitin synthase* gene with conserved alternative splice isoforms, which suggests that *chitin synthase 2* is a duplication specific to only certain beetle and fly lineages within the Holometabola [94].

A major characteristic of the milkweed bug is the distinctive red-orange and black aposematic (warning) coloration within the cuticle and epidermis that deters predators (*e.g.*, Figs. 1, 9, [20, 21]). For black coloration, the melanin synthesis pathway known from holometabolous insects (*e.g.*, [95, 96]) is conserved at the sequence (Fig. S5.15) and functional [97, 98] level in *Oncopeltus*, supporting conservation in hemimetabolous lineages as well. In contrast, production of the primary warning coloration, pteridine red erythropterin [99], has not been as extensively studied and remains an open avenue for hemimetabolous research. Pterin pigments are synthesized from GTP through a series of enzymatic reactions [100]. Thus far in *Oncopeltus* we could identify orthologs of *punch*, which encodes a GTP cyclohydrolase [101], and *sepia*, which is required for the synthesis of the red eye pigment drosopterin [102]. The bright red color of *Oncopeltus* eggs may in part reflect chemical protection transmitted parentally [103]. Thus, further identification of pigmentation genes will provide fitness indicators for maternal contributions to developmental success under natural conditions (*i.e.*, the presence of egg predators).

### Chemoreception and metabolism in relation to feeding biology

Aposematic pigmentation advertises the fact that toxins in the milkweed seed diet are incorporated into the insects themselves, a metabolic feat that was independently acquired in *Oncopeltus* and the monarch butterfly (*Danaus plexippus*), which shares this food source and body coloration [37, 104]. Moreover, given the fundamental differences between phytophagous, mucivorous, and hematophagous diets, we investigated to what extent differences in feeding ecology across hemipterans are represented in their chemoreceptor and metabolic enzyme repertoires.

**Table 2.** **Numbers of chemoreceptor genes/proteins per family in selected insect species.** In some cases the number of proteins is higher than the number of genes due to an unusual form of alternative splicing, which is particularly notable for the *Oncopeltus* GRs. Data are shown for four Hemiptera as well as *Drosophila melanogaster*, the body louse *Pediculus humanus*, and the termite *Zootermopsis nevadensis* [11, 12, 105-109].

| Species | Odorant | Gustatory | Ionotropic |
| --- | --- | --- | --- |
| *Oncopeltus fasciatus*<sup>1</sup> | 120/121 | 115/169 | 37/37 |
| *Cimex lectularius*<sup>1,2</sup> | 48/49 | 24/36 | 30/30 |
| *Rhodnius prolixus*<sup>1,2</sup> | 116/116 | 28/30 | 33/33 |
| *Acyrtosiphon pisum*<sup>3</sup> | 79/79 | 77/77 | 19/19 |
| *Pediculus humanus*<sup>2</sup> | 12/13 | 6/8 | 14/14 |
| *Zootermopsis nevadensis* | 70/70 | 87/90 | 150/150 |
| *Drosophila melanogaster* | 60/62 | 60/68 | 65/65 |

<sup>1</sup> Hemiptera: Heteroptera
<sup>2</sup> independent acquisitions of hematophagy [16]
<sup>3</sup> Hemiptera, phloem-feeding

Insects must smell and taste their environment to locate and identify food, mates, oviposition sites, and other essential cues. Perception of the enormous diversity of environmental chemicals is primarily mediated by the odorant (OR), gustatory (GR), and ionotropic (IR) families of chemoreceptors, which each encode tens to hundreds of distinct proteins [109-112]. Chemoreceptor family size appears to correlate with feeding ecology. *Oncopeltus* retains a moderate complement of each family, while species with derived fluid nutrition diets (sap or blood) have relatively depauperate OR and GR families (Table 2, Supplemental Note 5.3.f). In detail, a few conserved orthologs such as the OrCo protein and a fructose receptor are found across species, but other subfamilies are lineage specific. *Oncopeltus* and *Acyrthosiphon* retain a set of sugar receptors that was lost independently in the blood-feeding bugs (*Rhodnius* [11], *Cimex* [12]) and body louse (*Pediculus* [108]). Conversely, *Oncopeltus* and *Cimex* retain a set of candidate carbon dioxide receptors, a gene lineage lost from *Rhodnius*, *Acyrthosiphon*, and *Pediculus* [11, 12, 105], but which is similar to a GR subfamily expansion in the more distantly related hemimetabolous termite (Isoptera, [106]). Comparable numbers of IRs occur across the Heteroptera. In addition to a conserved set of orthologs primarily involved in sensing temperature and certain acids and amines, *Oncopeltus* has a minor expansion of IRs distantly related to those involved in taste in *Drosophila*. The major expansions in each insect lineage are the candidate “bitter” GRs ([113], Supplemental Note 5.3.f, Fig. S5.19). In summary, *Oncopeltus* exhibits moderate expansion of specific subfamilies likely to be involved in host plant recognition, consistent with it being a preferentially specialist feeder with a potentially patchy food source [21, 114].

As host plant recognition is only the first step, we further explored whether novel features of the *Oncopeltus* gene set are directly associated with its diet. We therefore used the CycADS annotation pipeline [115] to reconstruct the *Oncopeltus* metabolic network. The resulting BioCyc metabolism database for *Oncopeltus* (“OncfaCyc”) was then compared with those for 26 other insect species ([116], http://arthropodacyc.cycadsys.org/), including three other hemipterans: the pea aphid, the green peach aphid, and the kissing bug (Tables 3-4). For a global metabolism analysis, we detected the presence of 1085 Enzyme Commission (EC) annotated reactions with at least one protein in the *Oncopeltus* genome (Supplemental Note 6.4, Table S6.10). Among these, 10 enzyme classes (represented by 17 genes) are unique and 17 are missing when compared to the other insects (Table 4, Table S6.11).

We then specifically compared amino acid metabolism in the four hemipterans representing the three different diets. Eight enzymes are present only in *Oncopeltus* (Table 4), including the arginase that degrades arginine (Arg) into urea and ornithine, a precursor of proline (Pro). Given this difference, we extended our analysis to assess species’ repertoires for the entire urea cycle (Fig. 10a, Table S6.13). *Oncopeltus* and six other species can degrade Arg but cannot synthesize it (Fig. 10b). Only the other three hemipterans can neither synthesize nor degrade Arg via this cycle (Fig. 10c), while most species have an almost complete cycle (Fig. 10d). This suggests that the ability to synthesize Arg was lost at the base of the Hemiptera, with subsequent, independent loss of Arg degradation capacity in the aphid and *Rhodnius* lineages. Retention of Arg degradation in *Oncopeltus* might be linked to the milkweed seed food source, as most seeds are very rich in Arg [117], and Arg is indeed among the metabolites detected in *Oncopeltus* [118]. However, the monarch butterfly is one of only a handful of species that retains the complete Arg pathway (Fig. 10d: blue text). Despite a shared food source, these species may therefore differ in their overall Arg requirements, or – in light of a possible group benefit of *Oncopeltus* aggregation during feeding ([21]; *e.g.*, Fig. 1b) – in their efficiency of Arg uptake.

**Fig. 10.**
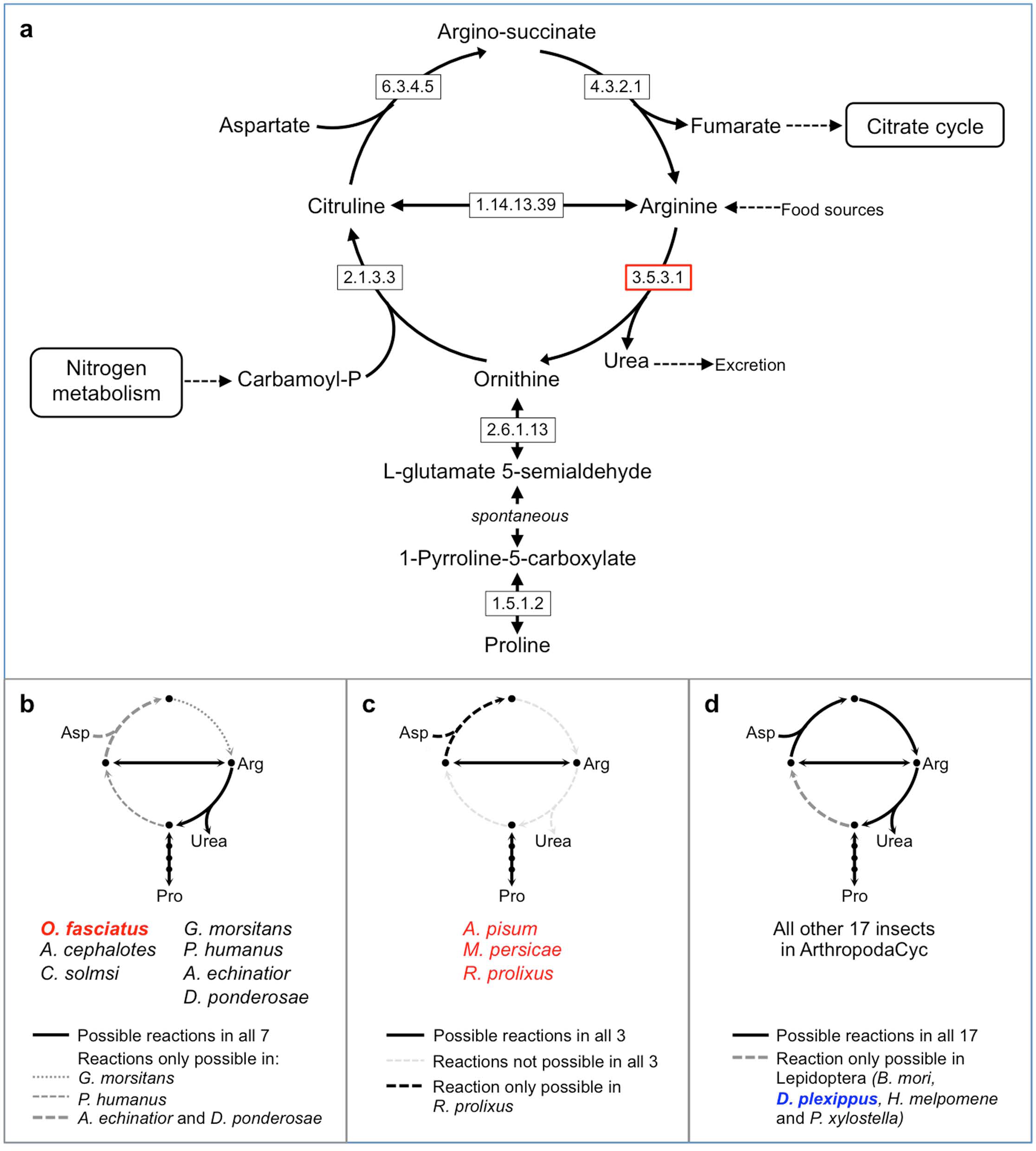
**Comparison of the urea cycle of *Oncopeltus* with 26 other insect species.** **(a)** Detailed diagram of the urea cycle (adapted from KEGG). **(b)** Group of 7 species, including *Oncopeltus*, for which Arg degradation via arginase (3.5.3.1), but not synthesis, is possible. **(c)** Group of 3 species for which neither the degradation nor synthesis of arginine via the urea cycle is possible (the three other hemipterans in this analysis). **(d)** Group of 17 species sharing a complete (or almost complete) urea cycle. Hemiptera are identified in red text and the milkweed-feeding monarch butterfly is in blue text. Enzyme names corresponding to EC numbers: 1.5.1.2 = pyrroline-5-carboxylate reductase; 1.14.13.39 = nitric-oxide synthase; 2.1.3.3 = ornithine carbamoyltransferase; 2.6.1.13 = ornithine aminotransferase; 3.5.3.1 = arginase; 4.3.2.1 = argininosuccinate lyase; 6.3.4.5 = argininosuccinate synthase. Analyses based on OGS v1.1.

Other enzymes are also present only in the milkweed bug compared to the other examined hemipterans (Table S6.12). Like other insects [116], *Oncopeltus* retains the ability to degrade tyrosine (Tyr). This pathway was uniquely lost in the aphids, where this essential amino acid is jointly synthesized – and consumed – by the aphid host and its endosymbiotic bacteria [6, 7, 17, 119]. Conversely, a gain specific to the milkweed bug lineage was the duplication of the Na+/K+ ATPase alpha subunits whose amino acid substitutions confer resistance to milkweed cardenolides [37, 120]. In the *Oncopeltus* genome, we find support for the recent nature of these duplications: the genes encoding subunits ATPα1B and ATPα1C occur as a tandem duplication, notably on a scaffold that also harbors one of the clustered ZF271-like gene expansions (see above).

**Table 3.** **Hemipteran ArthropodaCyc database summaries.** Overview statistics for the newly created database for *Oncopeltus fasciatus* (Ofas) in comparison with public databases for *Rhodnius prolixus* (Rpro), *Acyrthosiphon pisum* (Apis), and *Myzus persicae* (Mper) available from [116]. Based on OGS v1.1.

| Species ID<br>Gene set ID | *Ofas*<br>OGS v1.1 | *Rpro*<br>RproC1.1<br>(Built on<br>RproC1<br>assembly) | *Apis*<br>OGS v2.1b<br>(Built on<br>Acyr_2.0<br>assembly) | *Mper*<br>Clone G006<br>v1.0 | *Mper*<br>Clone O<br>v1.0 |
| --- | --- | --- | --- | --- | --- |
| CycADS Database ID | OncfaCyc | RhoprCyc | AcypiCyc<br>v2.1b | Myzpe_G006<br>Cyc | Myzpe_O<br>Cyc |
| Total mRNA<sup>1</sup> | 19,673 | 15,437 | 36,195 | 24,814 | 24,770 |
| Pathways | 294 | 312 | 307 | 319 | 306 |
| Enzymatic Reactions | 2,192 | 2,366 | 2,339 | 2,384 | 2,354 |
| Polypeptides | 19,820 | 15,471 | 36,228 | 24,849 | 24,805 |
| Enzymes | 3,050 | 2,660 | 5,087 | 4,646 | 4,453 |
| Compounds | 1,506 | 1,665 | 1,637 | 1,603 | 1,655 |

<sup>1</sup> In the BioCyc databases all splice variants are counted in the summary tables for genes.

**Table 4.** **Hemipteran ArthropodaCyc annotations of metabolic genes.** Taxonomic abbreviations are as in Table 3.

|  | *Ofas* | *Rpro* | *Apis* | *Mper* |
| --- | --- | --- | --- | --- |
| **Global metabolism** |  |  |  |  |
| EC<sup>1</sup> present in the genome | 1085 | 1241 | 1288 | 1222 |
| EC unique to this genome<sup>2</sup> | 10 | 13 | 23 | 5 |
| EC missing only in this genome<sup>2</sup> | 17<sup>4</sup> | 8 | 2 | 6 |
| **Amino acid metabolism (KEGG)** |  |  |  |  |
| EC present in the genome | 169 | 188 | 195 | 185 |
| EC unique to this genome<sup>2</sup> | 2 | 1 | 6 | 1 |
| EC missing only in this genome<sup>2</sup> | 5 | 2 | 0 | 2 |
| EC unique to this genome<sup>3</sup> | 8 | 10 | 12 | 8 |
| EC missing only in this genome<sup>3</sup> | 14 | 5 | 0 | 2 |

<sup>1</sup> "EC" refers to the number of proteins, as represented by their unique numerical designations within the Enzyme Commission (EC) classification system for enzymes and their catalytic reactions.
<sup>2</sup> in comparison to all other insects from ArthropodaCyc
<sup>3</sup> in comparison among the four hemipterans
<sup>4</sup> includes three EC categories added in OGS v1.2 (see also Table S6.11)

## Conclusions

The integrated genomic and transcriptomic resources presented here for the milkweed bug *Oncopeltus fasciatus* (Figs. 2,5) underpin a number of insights into evolutionary and developmental genomics. Our macroevolutionary comparisons across insect orders, now extended to the hemimetabolous Hemiptera, reveal unexpected patterns of molecular evolution. We also show how hemipteran feeding ecology and suites of related biological characters are reflected in the genome.

The gene structure trends we identified, with lineage predominating over genome size as a predictor and with many intron gains in the hemipteroid lineage (Figs. 6,7), offer initial parameters and hypotheses for the Hemiptera, Coleoptera, and Diptera. Such ordinal-level parameters can be evaluated against new species’ data and also inform customized pipelines for automated gene model predictions. At the same time, it will be interesting to explore the ramifications of hemipteroid intron gains, as there are few documented lineages with episodic intron gain [77]. For example, possessing more, small exons may provide greater scope to generate protein modularity via isoforms based on alternative exon usage [121]. Furthermore, with the larger genome sizes and lower gene densities of hemipteroids compared to the well-studied Hymenoptera, it remains open whether hemipteroid gene and intron size may also correlate with recombination rates [122].

Our analyses also highlight new directions for future experimental research, building on *Oncopeltus*’s long-standing history as a laboratory model and its active research community in the modern molecular genetics era (*e.g.*, Fig. 9, [25-27]). Functional testing will clarify the roles of genes we have identified as unique to the Hemiptera, including those implicated in chemical protection, bacterial and plant cell wall metabolism, or encoding wholly novel proteins (Figs. 3,8, see also Supplemental Note 2.2). Meanwhile, the prominent and species-specific expansions specifically of ZF271-like zinc fingers (Fig. 4), combined with the absence of the co-repressor KAP-1 in insects, argues for investigation into alternative interaction partners, which could clarify the nature of these zinc fingers’ regulatory roles and their binding targets.

One key output of this study is the generation of a metabolism database for *Oncopeltus*, contributing to the ArthropodaCyc collection (Table 3). In addition to comparisons with other species (Fig. 10), this database can also serve as a future reference for studies that use *Oncopeltus* as an ecotoxicology model (*e.g.*, [123]). While we have primarily focused on feeding ecology in terms of broad comparisons between phytophagy and fluid feeding, *Oncopeltus* is also poised to support future work on nuances among phytophagous species. Despite its milkweed diet in the wild, the lab strain of *Oncopeltus* has long been adapted to feed on sunflower seeds, demonstrating a latent capacity for more generalist phytophagy [114]. This potential may also be reflected in a larger gustatory receptor repertoire than would be expected for an obligate specialist feeder (Table 2). Thus, *Oncopeltus* can serve as a reference species for promiscuously phytophagous pests such as the stink bug. Finally, we have identified a number of key genes implicated in life history trade-offs in *Oncopeltus*, for traits such as cardenolide tolerance, pigmentation, and plasticity in reproduction under environmental variation. The genome data thus represent an important tool to elucidate the proximate mechanisms of fundamental aspects of life history evolution in both the laboratory and nature.

## Methods

(More information is available in the supplementary materials, Additional file 1.)

### Milkweed bug strain, rearing, and DNA/RNA extraction

The milkweed bug *Oncopeltus fasciatus* (Dallas), Carolina Biological Supply strain (Burlington, North Carolina, USA), was maintained in a laboratory colony under standard husbandry conditions (sunflower seed and water diet, 25 ºC, 12:12 light-dark photoperiod). Voucher specimens for an adult female (record # ZFMK-TIS-26324) and adult male (record # ZFMK-TIS-26325) have been preserved in ethanol and deposited in the Biobank of the Centre for Molecular Biodiversity Research, Zoological Research Museum Alexander Koenig, Bonn, Germany (https://www.zfmk.de/en/biobank).

Genomic DNA was isolated from individual, dissected adults using the Blood & Cell Culture DNA Midi Kit (G/100) (Qiagen Inc., Valencia, California, USA). Total RNA was isolated from individual, dissected adults and from pooled, mixed-instar nymphs with TRIzol Reagent (Invitrogen/Thermo Fisher Scientific, Waltham, Massachusetts, USA). Dissection improved accessibility of muscle tissue by disrupting the exoskeleton, and gut material was removed.

### Genome size calculations (flow cytometry, *k*-mer estimation)

Genome size estimations were obtained by flow cytometry with Hare and Johnston's protocol [124]. Four to five females and males each from the Carolina Biological Supply lab strain and a wild strain (collected from Athens, Georgia, USA; GPS coordinates: 33° 56' 52.8216'' N, 83° 22' 38.3484'’ W) were measured (see also Supplemental Note 2.1.a). At the bioinformatic level, we attempted to estimate genome size by *k*-mer spectrum distribution analysis for a range of *k*=15 to 34 counted with Jellyfish 2.1.4 [125] and bbmap [126], graphing these counts against the frequency of occurrence of *k*-mers (depth), and calculating genome size based on the coverage at the peak of the distribution (Supplemental Note 2.1.b).

### Genome sequencing, assembly, annotation, and official gene set overview

Library preparation, sequencing, assembly, and automatic gene annotation were conducted at the Baylor College of Medicine Human Genome Sequencing Center (as in [12, 30]). About 1.1 billion 100-bp paired-end reads generated on an Illumina HiSeq2000s machine were assembled using ALLPATHS-LG [127], from two paired-end (PE) and two mate pair (MP) libraries specifically designed for this algorithm (Supplemental Note 1). Three libraries were sequenced from an individual adult male (180- and 500-bp PE, 3-kb MP), with the fourth from an individual adult female (8-10-kb MP). The final assembly (see metrics in Table 1) has been deposited in GenBank (accession GCA_000696205.1).

Automated annotation of protein-coding genes was performed using a Maker 2.0 annotation pipeline [128] tuned specifically for arthropods (Supplemental Note 3). These gene predictions were used as the starting point for manual curation via the Apollo v.1.0.4 web browser interface [129], and automatic and manual curations were compiled to generate the OGS (see also Supplemental Note 4). Databases of the genome assembly, Maker automatic gene predictions, and OGS v1.1 are available through the i5K Workspace@NAL [130], and the Ag Data Commons data access system of the United States Department of Agriculture's (USDA) National Agricultural Library as individual citable databases [131-133]. The current version of the gene set, OGS v1.2, is deposited at GenBank as an ‘annotation-only’ update to the Whole Genome Shotgun project (accession JHQO00000000). Here, we describe version JHQO02000000. The annotations can be downloaded from NCBI’s ftp site, ftp://ftp.ncbi.nlm.nih.gov/genomes/all/GCA/000/696/205/GCA_000696205.1_Ofas_1.0/.

### Repeat content analysis

Repetitive regions were identified in the *Oncopeltus* genome assembly with RepeatModeler Open-1.0.8 [134] based on a species-specific repeat library generated *de novo* with RECON [135], RepeatScout [136], and Tandem Repeats Finder [137]. Then, RepeatMasker Open-4.0 [138] was used to mask repeat sequences based on the RepeatModeler library. Given the fragmented nature of the assembly, we attempted to fill and close assembly gaps by sequencing additional material, generating long reads with single molecule real time sequencing on a PacBio RS II machine (estimated coverage of 8x). Gap filling on the Illumina assembly scaffolds was performed with PBJelly version 13.10.22, and the resulting assembly was used for repeat content estimation and comparison with *Cimex lectularius* and *Acyrthosiphon pisum* (see also Supplemental Note 2.3).

### Transcriptome resources

Total RNA from three distinct life history samples (pooled, mixed-instar nymphs; an adult male; an adult female) was also sequenced on an Illumina HiSeq2000s machine, producing a total of 72 million 100-bp paired-end reads (Supplemental Note 1.3, Table S1.1; GenBank Bioproject: PRJNA275739). These expression data were used to support the generation of the OGS at different stages of the project: as input for the evidence-guided automated annotation with Maker 2.0 (Supplemental Note 3), as expression evidence tracks in the Apollo browser to support the community curation of the OGS, and, once assembled into a *de novo* transcriptome, as a point of comparison for quality control of the OGS.

The raw RNA-seq reads were pre-processed by filtering out low quality bases (phred score <30) and Truseq adapters with Trimmomatic-0.30. Further filtering removed ribosomal and mitochondrial RNA sequences with Bowtie 2 [139], based on a custom library built with all hemipteran ribosomal and mitochondrial RNA accessions from NCBI as of 7^th^ February 2014 (6,069 accessions). The pooled, filtered reads were mapped to the genome assembly with Tophat2-PE on CyVerse [140]. A second set of RNA-seq reads from an earlier study (“published adult” dataset, [37]) was also filtered and mapped in the same fashion, and both datasets were loaded into the *Oncopeltus* Apollo instance as evidence tracks (under the track names “pooled RNA-seq - cleaned reads” and “RNA-seq raw PE reads Andolfatto et al”, respectively).

Additionally, a *de novo* transcriptome was generated from our filtered RNA-seq reads (pooled from all three samples prepared in this study) using Trinity [141] and TransDecoder [142] with default parameters. This transcriptome is referred to as “i5K”, to distinguish it from a previously published maternal and early embryonic transcriptome for *Oncopeltus* (referred to as “454”, [36]). Both the i5K and 454 transcriptomes were mapped to the genome assembly with GMAP v. 2014-05-15 on CyVerse. These datasets were also loaded into the Apollo browser as evidence tracks to assist in manual curation.

### Life history stage-specific and sex-specific expression analyses in hemipteroids

Transcript expression of the OGS v1.1 genes was estimated by running RSEM2 [143] on the filtered RNA-seq datasets for the three i5K postembryonic stages against the OGS v1.1 cDNA dataset. Transcript expression was then based on the transcripts per million (TPM) value. The TPM values were processed by adding a value of 1 (to avoid zeros) and then performing a log2-transformation. The number of expressed genes per RNA-seq library was compared for TPM cutoffs of >1, >0.5, and >0.25. A >0.25 cutoff was chosen, which reduced the number of expressed genes by 6.6% compared to a preliminary analysis based on a simple cutoff of ≥10 mapped reads per transcript, while the other TPM cutoffs were deemed too restrictive (reducing the expressed gene set by >10%). This analysis was also applied to the “published adult” dataset [37]. To include embryonic stages in the comparison, transcripts from the 454 transcriptome were used as blastn queries against the OGS v1.1 cDNA dataset (cutoff e-value <10^-5^). The results from all datasets were converted to binary format to generate Venn diagrams (Fig. 2b).

Statistically significant sex-specific and developmental stage-specific gene enrichment was determined from RNA-seq datasets according to published methods [144, 145], with modifications. Data from *Oncopeltus* (see previous methods section, Bioproject: PRJNA275739) were compared between stages and pairwise with the hemipterans *Cimex lectularius*, PRJNA275741; *Acyrthosiphon pisum*, PRJNA209321; and *Pachypsylla venusta*; PRJNA275248; as well as with the hemipteroid *Frankliniella occidentalis* (Thysanoptera), PRJNA203209 (see also Fig. 2c, Supplemental Note 2.4).

### Protein gene orthology assessments via OrthoDB and BUSCO analyses

These analyses follow previously described approaches and with the current database and pipeline versions [1, 43, 45, 146]. See Supplemental Note 6.1 for further details.

### Global transcription factor identification

Likely transcription factors (TFs) were identified by scanning the amino acid sequences of predicted protein-coding genes for putative DNA binding domains (DBDs), and when possible, the DNA binding specificity of each TF was predicted using established procedures [58]. Briefly, all protein sequences were scanned for putative DBDs using the 81 Pfam [147] models listed in Weirauch and Hughes [148] and the HMMER tool [149], with the recommended detection thresholds of Per-sequence Eval < 0.01 and Per-domain conditional Eval < 0.01. Each protein was classified into a family based on its DBDs and their order in the protein sequence (*e.g*., bZIPx1, AP2x2, Homeodomain+Pou). The resulting DBD amino acid sequences were then aligned within each family using Clustal Omega [150], with default settings. For protein pairs with multiple DBDs, each DBD was aligned separately. From these alignments, the sequence identity was calculated for all DBD sequence pairs (*i.e*., the percent of amino acid residues that are identical across all positions in the alignment). Using previously established sequence identity thresholds for each family [58], the predicted DNA binding specificities were mapped by simple transfer. For example, the DBD of OFAS001246-RA is 98% identical to the *Drosophila melanogaster* Bric a Brac 1 (Bab1) protein. Since the DNA binding specificity of Bab1 has already been experimentally determined, and the cutoff for the Pipsqueak family TFs is 85%, we can infer that OFAS001246-RA will have the same binding specificity as *Drosophila* Bab1.

### RNA interference

Double-stranded RNA (dsRNA) was designed to target the final, unique exon of the *broad* isoforms *Z2, Z3*, and *Z4*. A portion of the coding sequence for the zinc finger region from these exons (179 bp, 206 bp, and 216 bp, respectively) was cloned into a plasmid vector and used as template for *in vitro* RNA synthesis, using the gene-specific primer pairs: Of-Z2_fwd: 5′-ATGTGGCAGACAAGCATGCT-3′; Of-Z2_rev: 5′-CTAAAATTTGACATCAGTAGGC-3′; Of-Z3_fwd: 5′-ccttctcctgttactactcac-3′; Of-Z3_rev: 5′-ttatatgggcggctgtccaa-3′; Of-Z4_fwd: 5′-AACACTGACCTTGGTTACACA-3′; Of-Z4_rev: 5′-TAGGTGGAGGATTGCTAAAATT-3′. Two separate transcription reactions (one for each strand) were performed using the Ambion MEGAscript kit (Ambion, Austin, Texas, USA). The reactions were purified by phenol/chloroform extraction followed by precipitation as described in the MEGAscript protocol. The separate strands were re-annealed in a thermocycler as described previously [33]. Nymphs were injected with a Hamilton syringe fitted with a 32-gauge needle as described [55]. The concentration of *Of-Z2*, *Of-Z3* and *Of-Z4* dsRNA was 740 ng/µl, 1400 ng/µl, and 1200 ng/µl, respectively. All nymphs were injected within 8 hours of the molt to the fourth (penultimate juvenile) instar (n ≥12 per treatment: see Fig. 9). Fore- and hindwings were then dissected from adults and photographed at the same scale as wings from wild type, uninjected controls.

### CycADS annotation and OncfaCyc database generation

We used the Cyc Annotation Database System (CycADS, [115]), an automated annotation management system, to integrate protein annotations from different sources into a Cyc metabolic networks reconstruction that was integrated into the ArthropodaCyc database. Using our CycADS pipeline, *Oncopeltus fasciatus* proteins from the official gene set OGS v1.1 were annotated using different methods – including KAAS [151], PRIAM [152], Blast2GO [153, 154], and InterProScan with several approaches [155] – to obtain EC and GO numbers. All annotation information data were collected in the CycADS SQL database and automatically extracted to generate appropriate input files to build or update BioCyc databases [156] using the Pathway Tools software [157]. The OncfaCyc database, representing the metabolic protein-coding genes of *Oncopeltus*, was thus generated and is now included in the ArthropodaCyc database, a collection of arthropod metabolic network databases ([116], http://arthropodacyc.cycadsys.org/).

## Additional Files

Additional file 1: Supplementary figures, tables, methods, and other text. (PDF)

Additional file 2: Large supporting tables. (XLSX)

Additional file 3: Chemoreceptor sequences in FASTA format. (TXT)

## Acknowledgements

We thank Dorith Rotenberg (Kansas State University, currently North Carolina State University, USA) and Michael Sparks (Agricultural Research Service, United States Department of Agriculture, USA) for generously making available the unpublished genome assemblies of the fellow hemipteroid i5K species *Frankliniella occidentalis* and *Halyomorpha halys*, respectively, for use in specific analyses presented here. Similarly, we thank Hans Kelstrup and Lynn Riddiford (Janelia Farm Research Campus, HHMI, USA) for sharing unpublished data on *Of-E75A* RNAi. We thank George Coupland (Max Planck Institute for Plant Breeding Research, Cologne, Germany) as well as Lisa Czaja, Kurt Steuber, and Bruno Huettel (Max Planck Genome Centre Cologne, Germany) for conducting the PacBio sequencing and providing support with data handling. We also thank Oliver Niehuis (Albert Ludwig University, Freiburg, Germany) and Alexander Klassmann (University of Cologne, Germany) for discussions on *k*-mer and gene structure analyses, respectively, Sarah Kingan (University of Rochester, USA) for assistance with LGT phylogenies, as well as Jeanne Wilbrandt (Zoologisches Forschungsmuseum Alexander Koenig, Bonn, Germany) for comments on the manuscript.

## Availability of Data and Materials

All sequence data are publically available at the NCBI, bioproject number PRJNA229125. In addition, gene models and a browser are available at the National Agricultural Library ([131-133], https://i5k.nal.usda.gov/Oncopeltus_fasciatus).

## Authors’ Contributions

KAP and StR conceived the project. KAP managed and coordinated the project. KAP and SK provided specimens for sequencing and performed DNA and RNA extractions. StR, SD, SLL, HC, HVD, HD, YH, JQ, SCM, DSTH, KCW, DMM, and RAG constructed libraries and performed sequencing. StR, SCM, and DSTH performed the genome assembly and automated gene prediction. IMVJ, JSJ, and PJM analyzed genome size. IMVJ, VK, PH, and KAP contributed to repetitive content analyses. AD, RR, JHW, KAP, and SK performed bacterial scaffold detection and LGT analyses. MCMT developed Apollo software. KAP, IMVJ, MCMT, CPC, C-YL, and MFP implemented Apollo-based manual curation. KAP, IMVJ, JBB, DE, YS, HMR, DA, CGCJ, BMIV, EJD, CSB, C-CC, Y-TC, ADC, AGC, AJJC, PKD, EMD, CGE, MF, NG, TH, Y-MH, ECJ, TEJ, JWJ, AK, ML, MRL, H-LL, YL, SRP, LP, MP, PNR, RRP, SiR, LS, MES, JS, ES, JNS, OT, LT, MVDZ, SV, and AJR participated in manual curation and contributed to the Supplemental Notes. IMVJ, KAP, DSTH, M-JMC, CPC, C-YL, and MFP performed curation quality control and generated the OGS. IMVJ, KAP, CJH, and JBB generated *de novo* transcriptomes and performed life history stage expression analyses. RMW, PI, KAP, and EMZ performed orthology and phylogenomic analyses. MTW, KAP, IMVJ, PH, and BMIV performed transcription factor analyses. EJD conducted analyses of DNA methylation. KAP, PH, and RJS contributed to comparative analyses of gene structure. DE conducted the RNAi experiments. SC, PB-P, GF, and NP generated and performed comparative analyses on the OncfaCyc database. KAP, IMVJ, JBB, DE, YS, SC, HMR, and MTW wrote the manuscript. KAP, IMVJ, JBB, DE, YS, SC, HMR, MFP, RMW, PI, MTW, StR, PJM, and AK edited the manuscript. IMVJ and KAP organized the Supplementary Materials. All authors approved the final manuscript.

## Ethics Approval and Consent to Participate

Not applicable.

